# Scalable 3D cell-interaction analysis via supercell graphs for prostate cancer risk stratification

**DOI:** 10.64898/2026.07.07.736891

**Authors:** Yujie Zhao, Sarah S.L. Chow, Renao Yan, David Brenes, Robert Serafin, Cristina Almagro-Pérez, Andrew H Song, Priti Lal, Emily Chan, Michelle Downes, Elena Baraznenok, Jennifer Salguero Lopez, Anant Madabhush, Faisal Mahmood, Lawrence D. True, Jonathan T.C. Liu

## Abstract

Cellular interactions underlie fundamental biological processes but are not fully represented in conventional 2D histology images. While 3D pathology allows for more-accurate construction of cell-level graphs, machine-learning models are computationally unwieldy and prone to overfitting, especially when dealing with small cohorts. Here, we introduce **SCALE3D**, a **S**uper**C**ell graph **A**nalysis framework for **L**arg**E 3D** pathology datasets. In SCALE3D, spatially adjacent and morphologically similar cells are grouped into functional “supercells.” Supercell subtypes are defined via morphology-based clustering and 3D graphs connecting these supercells are used to model their interactions. Validation was performed with 76 radical prostatectomy specimens from patients with known 5-year biochemical recurrence (BCR) outcomes. SCALE3D-derived features achieve higher performance for BCR prediction than established 3D nuclear and glandular morphological features. Combining these complementary features further improves prediction performance. Compared to individual cell-level 3D graphs, SCALE3D maintains comparable prognostic performance with improved noise tolerance while reducing computational times by up to 1,000-fold.

## Introduction

Prostate cancer (PCa) is the most commonly diagnosed cancer and the second leading cause of cancer-related deaths among men^1^. In current clinical practice, histopathological evaluation of thin hematoxylin and eosin (H&E)-stained tissue (∼5 µm) remains the gold standard for prostate cancer diagnosis and risk stratification. When prostate cancer is identified in conventional 2D H&E histology slides, the accurate grading of the cancer is critical since most prostate cancers are relatively indolent and do not require aggressive treatment. Standard-of-care Gleason grading of prostate cancers, which is one of the most important factors influencing treatment decisions, is based entirely on interpreting the glandular morphology seen in H&E histology slides^2^. However, studies have shown that other tissue structures are prognostic as well, such as nuclear morphology^3^. Beyond these morphological features, increasing evidence suggests that the spatial organization and interaction between different cell types is also correlated with disease aggressiveness^4,5^. For example, immune-excluded tumor microenvironments are associated with tumor progression^4^ whereas the presence of intratumoral tertiary lymphoid structures correlates with lower risk of progression^5^.

Most prior studies rely on 2D pathological images, which sample only a small fraction of the specimen (typically ≤ 1%), leading to incomplete and ambiguous representations of complex, heterogeneous 3D tissue architectures and cellular interactions^6^. In contrast, recent advances in high-throughput, non-destructive 3D pathology have enabled comprehensive sampling of large tissue volumes. Among the various approaches to 3D pathology^7–12^, open-top light-sheet (OTLS) microscopy can generate high-quality 3D datasets with spatial resolution comparable to conventional 2D histology and with contrast that mimics the appearance of H&E staining^6,7,11,13,14^. As described in recent protocols, H&E-analog staining can be performed rapidly and inexpensively in thick tissue using small-molecule fluorophores, followed by optical clearing to enable deep 3D microscopy^14,15^. Compared with conventional 2D histology, 3D pathology provides a framework to capture the 3D spatial organization of the tumor microenvironment and to evaluate its prognostic value (**Fig. 1a**)^16,17^.

**Figure 1.**
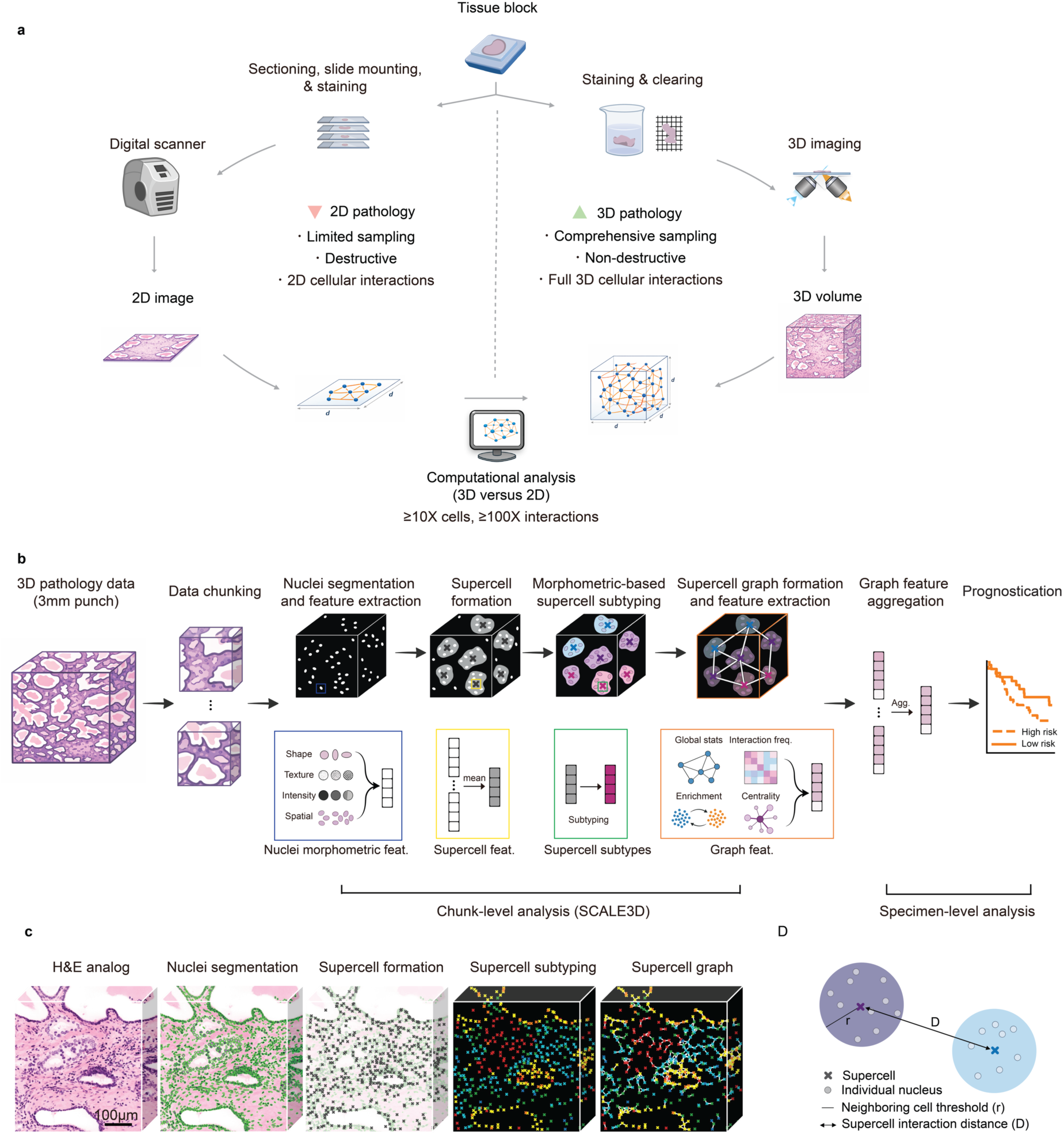
(a) Conventional 2D histopathology captures only a limited set of cellular interactions in a few thin sections cut from one arbitrary side of a specimen, whereas 3D pathology preserves all cellular interactions within a much larger tissue volume, thereby enabling a more comprehensive and accurate evaluation of these heterogeneous microenvironments. However, the large number of cells and interaction pairs in 3D pathology datasets poses a computational challenge that motivates the development of biologically informed strategies to create a coarse-grained representation of the cell interactions. (b) SCALE3D overview. 3D pathology datasets are divided into smaller chunks for chunk-level analysis. Within each chunk, nuclei are segmented and represented by single-cell features (shape, texture, intensity, and spatial descriptors). Morphologically similar neighboring nuclei are grouped into supercells, generating supercell-level features. Supercells are then categorized into morphology-based subtypes and connected to form spatial graphs. From each supercell graph, graph-level features are extracted, including interaction frequencies, neighborhood enrichment, centrality, and global graph statistics. These chunk-level features are subsequently aggregated to produce specimen-level representations for prognostic modeling. (c) Image examples along the SCALE3D pipeline. Representative example of an H&E-analog image, nuclear segmentation, supercell formation, supercell subtyping (color indicates different supercell subtypes), and supercell graph. (d) Two key parameters in the SCALE3D pipeline. The neighboring cell threshold, r, defines the maximum distance within which neighboring nuclei may be grouped into the same supercell. The supercell interaction distance, D, defines the maximum distance at which two supercells may be connected (i.e. allowed to interact as an edge within a graph).

Cellular interactions are typically modeled by first performing nuclear segmentation, cell-type annotation, and representing tissues as cell graphs, in which cells are represented as nodes and their interactions as edges. However, large-scale graphs pose significant computational challenges for downstream analysis using either traditional machine learning or deep learning methods. Scaling graph analyses to 3D data further exacerbates these computational challenges – a single biopsy can generate 10–100 GB of high-resolution 3D pathology data with millions of cells and orders-of-magnitude more cell interactions. With such large datasets and limited specimens that are labeled/annotated, 3D graph-based models that operate at the individual-cell level are prone to overfitting with high levels of noise and poor computational efficiency, which limits translation to clinical settings^18–20^.

Prior studies have attempted to reduce the computational complexity of cell-graph analysis in various ways, such as analyzing a sub-population of representative cells^21,22^, aggregating features at the patch level^23^, or restricting analyses to predefined regions of interest^24^. Another data-compression approach is to group pixels/patches from homogeneous tissue regions into “superpixels” or “superpatches,” thereby enabling models to operate on fewer, more stable units^25–28^. These strategies can improve efficiency and reduce noise, providing smoother, more robust inputs for downstream analysis. However, these methods don’t preserve cells as the fundamental units, they may obscure cell morphology and are less well suited for modeling cell interactions.

From a biological perspective, individual cells rarely act in isolation. Instead, similar cells typically work collectively within spatially coherent neighborhoods to exert meaningful region-level biological functions^29–33^. Based on this general principle, we developed **SCALE3D**: a **S**uper**C**ell graph **A**nalysis framework for **L**arg**E 3D** pathology datasets. Rather than constructing graphs at the individual-cell level, SCALE3D aggregates spatially adjacent and morphologically similar nuclei into biologically functional “supercells”, leveraging a computationally efficient coarse-grained representation while preserving key aspects of tissue architecture and cellular context. Supercell subtypes are then defined via morphology-based clustering, with supercell spatial interactions quantified by extracting features derived from 3D radius graphs.

We applied SCALE3D to 3D pathology datasets (generated by OTLS microscopy) of 76 radical prostatectomy (RP) specimens from patients with known biochemical recurrence (BCR) outcomes, showing that SCALE3D features complement previously studied 3D morphological features for prostate cancer risk stratification. In addition, compared with standard cell-level graphs, SCALE3D demonstrates comparable prognostic performance and increased robustness to feature noise while greatly reducing computation times, underscoring the benefits of biologically informed graph compression for large 3D datasets with small cohorts. An overview of the study design, SCALE3D workflow, and major findings is provided in **Extended Data Video 1**.

## Results

### Overview of the pipeline

Although 3D pathology enables more accurate characterization of cellular interactions than 2D pathology (**Fig. 1a**), it introduces substantial scaling challenges. In our cohort, 3D specimens contain ≥10× more nuclei and ≥100× more candidate interactions (radius-based graph) than corresponding 2D sections (**Extended Data Fig. 1a**). This increase in graph complexity not only raises computational cost but also increases the risk of overfitting, particularly considering that cohort sizes tend to be limited in the nascent but growing field of 3D pathology.

To overcome these limitations, here we introduce the 3D supercell graph framework (SCALE3D, **Fig. 1b; Extended Data Video 2**). Our method leverages the general biological concept that cells rarely exert discernible functions in isolation, but rather operate through collective and coordinated cell interactions within the tissue microenvironment to have a palpable effect at the regional or organ level^29–33^. By adopting this concept, SCALE3D groups morphologically similar neighboring nuclei into biologically meaningful “supercells”. This design preserves essential biological information while providing a data-efficient representation for modeling large-scale 3D cellular interactions and evaluating their prognostic significance.

Given the computational demands of whole-specimen volumetric processing, each specimen is first partitioned into non-overlapping 3D chunks that are processed independently. The chunk-level pipeline is comprised of four main stages: (i) 3D nuclei segmentation, (ii) supercell formation, (iii) morphology-based supercell subtyping, and (iv) 3D supercell graph formation. For specimen-level analysis, chunk-level graph-derived feature vectors are aggregated to generate specimen-level representations for downstream clinical-outcome predictions (**Fig. 1b**; Methods).

For chunk-level analysis, individual nuclei are first segmented in 3D using Cellpose^34^ (**Extended Data Fig. 1b**). From the resulting nuclear masks, a comprehensive panel of 3D nuclear morphological features is extracted encompassing shape, intensity statistics, texture, and local spatial context (**Extended Data Table 1**). To maintain biologically relevant compartmentalization, nuclei are first stratified into epithelial and stromal regions using precomputed glandular masks^35^. Supercell formation and subtyping are then performed independently within each tissue region.

Supercells are generated using an iterative region-growing procedure. Starting from a randomly selected nucleus, neighboring nuclei within a radius r (neighboring-cell threshold) are evaluated as candidate supercell members. The nuclei are grouped together within a supercell only if their standardized morphometric feature vector (**Extended Data Table 1**) shows a high cosine similarity to the current supercell-level feature vector, which is defined as the running mean of the morphological feature vector of the nuclei already assigned to that supercell. The spatial centroid of each supercell is defined as the spatial average of its constituent nuclei centroids. This process is repeated until every nucleus in every 3D chunk is either assigned to a supercell or considered during the algorithm. All 3D chunks are processed in this manner. Supercell subtypes are then identified by applying Leiden clustering to the pooled supercell morphological feature vectors across chunks and specimens^36–38^ (Methods).

To model cellular interactions between spatially proximal supercells, we construct 3D radius graphs in which supercells are represented as nodes and the interactions between supercell pairs are represented as edges, defined within a tunable physical interaction distance **D**. Graphs are generated independently within each chunk. Graph-based features are then calculated to summarize how frequently each supercell subtype contacts or co-localizes with other supercell subtypes (**Extended Data Table 2**). Representative image examples along the SCALE3D pipeline are provided in **Fig. 1c**, including H&E-analog images, nuclear segmentation, supercell formation, supercell subtyping, and supercell graphs. An image atlas of example 2D cross sections from all specimens is provided in **Extended Data Fig. 2**.

Two key distance parameters, **r** (10–30 µm) and **D** (10–60 µm), were systematically varied in an ablation study to optimize the prognostic performance of SCALE3D features (**Fig. 1d).** These ranges were informed by prior multiplexed-imaging and cell-graph studies reporting that cells typically communicate and interact within a proximity of ∼20–40 µm^39–42^. In this study, these ranges were expanded to accommodate for the coarse-grained nature of the supercells.

### Optimization of the SCALE3D framework for risk stratification

The SCALE3D pipeline was applied to whole-volume 3D pathology datasets from a cohort of 76 archived FFPE radical prostatectomy specimens, with one specimen per patient. All patients had known 5-year biochemical recurrence (BCR) outcomes (i.e. whether patients had BCR within 5 years of surgery), with 44 patients without BCR (“non-BCR” group) and 32 patients with BCR (“BCR” group). The specimens imaged with 3D pathology consisted of 3-mm diameter punches extracted from prostatectomy regions that were deemed to represent the index lesion. To optimize the SCALE3D framework, we evaluated its predictive performance for binary 5-year BCR outcomes using Least Absolute Shrinkage and Selection Operator (LASSO) regularized logistic regression. Performance was quantified by the area under the receiver operating characteristic curve (AUC) when employing leave-one-out cross-validation (LOOCV).

Systematic parameter ablation was performed on the neighboring-cell threshold (r = 10–30 µm) and supercell interaction distance (D = 10–60 µm). Intermediate values yielded the highest performance across the experimental matrix of parameters (r ≈ 15 µm for the neighboring-cell threshold; D ≈ 20 µm for the supercell interaction distance), providing a balance between data compression and preservation of prognostic spatial interactions as indicated by classifier performance (**Fig. 2a)**.

**Figure 2.**
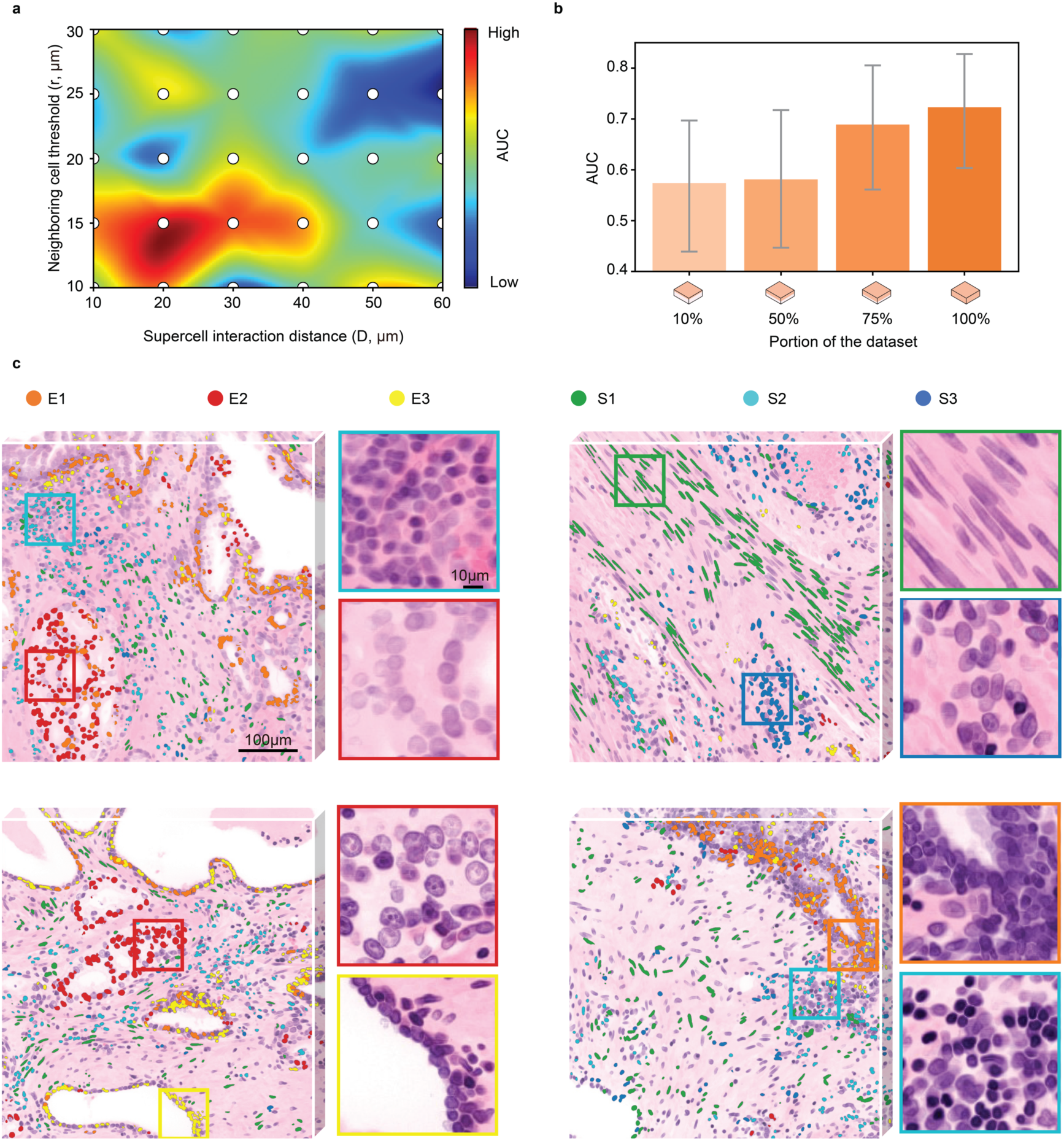
Optimization of the SCALE3D framework and morphometric characterization of supercell subtypes. (a) Parameter optimization. Heatmap showing cross-validated AUC across different combinations of neighboring-cell thresholds, r, and supercell interaction distances, D. Warmer colors indicate higher predictive performance. (b) 3D data contribution analysis. AUC values obtained using different proportions of the dataset, 10%, 50%, 75%, and 100%. (c) Morphometric characterization of E1-E3 (orange, red, yellow) and S1-S3 (green, cyan, blue). Nuclei are colored according to their assigned supercells, with representative zoomed-in regions shown for areas enriched for each subtype.

To assess the contribution of volumetric context, we evaluated model performance using progressively larger fractions of the 3D tissue thickness starting from the top surface (10%, 50%, 75%, and 100%). Predictive performance increased with greater volumetric coverage, and models using the full-thickness volumes achieved the highest AUC (**Fig. 2b**). These findings indicate that incorporating more 3D spatial context from a FFPE punch specimen enhances prognostic discrimination, re-affirming previous findings that also found this to be true for similar biopsy-sized specimens ^16,17^.

### Morphometric characterization of supercell subtypes

With the best supercell formation parameter value (r = 15 µm), morphometry-based clustering identified six distinct supercell subtypes, comprising three epithelial (E1–E3) and three stromal (S1–S3) subtypes. To qualitatively characterize these subtypes in a rough way, we show representative patches for each subtype in **Figure 2c**, and additional examples of representative image patches for each subtype in **Extended Data Figure 3**.

For epithelial clusters, E1 (orange) is characterized by well-formed glandular epithelial structures with variable nucleolar prominence. Pathologist reviewers note that E1 is seen in a range of glands from Gleason pattern 3-like carcinoma to benign atrophic glands. E2 (red) represents a mixture of carcinoma cells from Gleason pattern 3 and poorly formed gland variants of Gleason pattern 4, characterized by glands with irregular shapes and distorted or absent lumina. E3 (yellow) corresponds predominantly to benign glands with relatively small nuclei, inconspicuous nucleoli, and micropapillary structures. Adjacent basal cells peripheral to the luminal epithelial cells are variably present.

For stromal clusters, S1 (green) is composed of elongated spindle-shaped cells with small nuclei that do not form distinct architectural elements, consistent with fibromuscular stromal cells, including smooth muscle cells, myofibroblasts, and fibroblasts. S2 (cyan) consists of small, irregularly clustered cells with round nuclei and dense heterochromatin, morphologically consistent with lymphoid cells. S3 (blue) exhibits mixed morphologic characteristics of S1 and S2, with vaguely clustered architecture that is not obviously glandular.

### Evaluating the prognostic value of SCALE3D features

We evaluated the prognostic value of the SCALE3D features introduced in this study relative to established features such as 3D glandular^43^ and nuclear^44^ morphologies. Glandular morphological features were extracted at the specimen level, whereas nuclear morphological features were computed at the cell level and subsequently aggregated to the specimen level. In total, we extracted and examined 522 nuclear features, 39 glandular features, and 320 SCALE3D features. To ensure fair comparison across feature sets with different dimensionalities, feature selection was first performed within each cross-validation training fold, and top 20 features were used to train all classifiers. We compared the performance of these features individually and in combination. LASSO-regularized logistic regression was used for binary 5-year BCR classification and ridge-penalized Cox proportional hazards regression was performed for time-to-BCR analysis (Methods).

Single-modality performance: for binary classification, SCALE3D features alone achieved the highest AUC (0.723; 95% CI 0.60–0.83), outperforming glandular features (AUC 0.621; 95% CI 0.47–0.75) and nuclear features (AUC 0.638; 95% CI 0.51–0.75) (**Fig. 3a**). Kaplan–Meier analysis confirmed significant risk stratification for all modalities (nuclei: p = 0.0044, HR = 2.77, C-index = 0.66; gland: p = 0.0227, HR = 2.33, C-index = 0.62; SCALE3D: p = 0.00288, HR = 2.90, C-index = 0.65). These results establish that SCALE3D-derived features provide prognostic information comparable to or exceeding the established 3D nuclear and glandular features.

**Figure 3.**
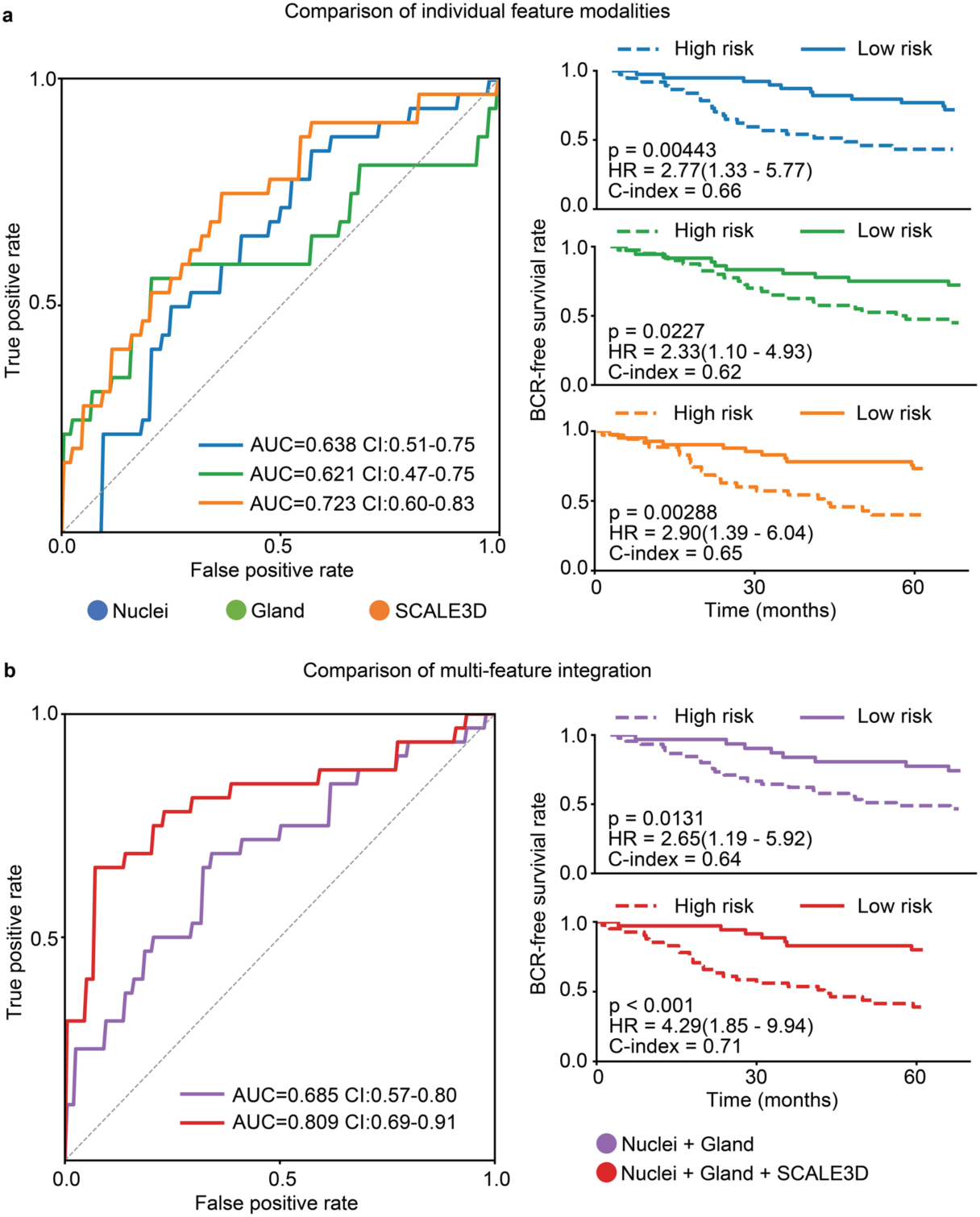
Prognostic evaluation of SCALE3D features. (a) Comparison of individual feature modalities. Left: ROC curves for nuclei-derived, gland-derived, and SCALE3D-derived features. Right: Kaplan–Meier curves showing risk stratification for BCR-free survival based on each feature modality. Each color indicates a distinct feature modality. (b) Comparison of multi-feature integration. Left: ROC curves for nuclei + gland features versus nuclei + gland +SCALE3D features. Right: Kaplan–Meier curves are shown for BCR-free survival corresponding to the integrated models. Each color indicates a distinct feature modality.

Multi-modality integration: we evaluated whether combining SCALE3D features with glandular and nuclear features further improved prognostic performance. Integrating SCALE3D features with nuclear and glandular morphological features yielded a superior AUC of 0.809 (95% CI 0.69–0.91), compared with 0.685 (95% CI 0.57–0.80) for nuclear + glandular features alone (**Fig. 3b**). Kaplan–Meier curves also showed enhanced separation with the full model (nuclei + gland + SCALE3D: p < 0.001, HR = 4.29, C-index = 0.71) than the reduced model (nuclei + gland: p = 0.0131, HR = 2.65, C-index = 0.64). To quantify the incremental value of SCALE3D features, we compared out-of-fold risk scores from reduced (nuclei + gland) versus full (nuclei + gland + SCALE3D) Cox models using likelihood ratio testing. Incorporating SCALE3D-derived features significantly improved model fit (χ^2^(1) = 8.45, p = 0.0036), indicating that SCALE3D features provide complementary prognostic information to established features. Due to the relatively small cohort size, we also performed permutation testing (randomly shuffling the outcome labels) to assess whether the observed predictive performance could arise by chance. Across 500 random label permutations, the mean AUC was 0.49 ± 0.11, and none exceeded the observed AUC of 0.809 (empirical p = 0.002), supporting the statistical validity of the full model.

To evaluate feature robustness, we assessed selection stability across the 76 LOOCV folds, defining selection frequency as the proportion of folds in which a feature was retained after feature preselection and LASSO shrinkage. Ten features demonstrated high stability (selection frequency ≥ 90%), indicating these features were consistently selected despite variations in the training data. Notably, six of these ten stable features were SCALE3D features. This preferential selection suggests that SCALE3D features capture reproducible and informative signals for BCR prediction, while their consistent selection across folds supports their robustness to sampling variability and reduces the likelihood that they arise from overfitting.

Among the stable SCALE3D features, epithelial–epithelial interaction variability emerged as an important signal: the standard deviation of enrichment Z-scores for epithelial subtype interactions was significantly elevated in the BCR group (**Extended Data Fig. 4a**), indicating increased heterogeneity in epithelial interaction patterns. In addition, epithelial–stromal interaction features showed distinct trends between groups with the mean enrichment Z-scores for E2 and S3 being higher in the non-BCR group, and the standard deviation of enrichment Z-scores for E2 and S3 being higher in the BCR group (**Extended Data Fig. 4b**). Finally, the median closeness centrality of E3 was significantly lower in the BCR group, suggesting that E3 subtypes were more spatially separated from other subtypes in recurrent cases (**Extended Data Fig. 4c; Extended Data Table 3**).

### Supercell versus individual cell-level graph analysis

Although the SCALE3D framework demonstrates the ability to capture prognostic information and is computationally advantageous, a concern is that the coarse-grained supercell representation may sacrifice biologically meaningful details and compromise downstream analytical performance. To this end, we compared supercell-level (SCALE3D) with a standard cell-level graph analysis. Due to computational constraints, this comparison was performed on a subset of the dataset, comprising 38 specimens and 25% of each 3D pathology volume.

For both representations, we applied identical processing steps, including morphology-based subtyping via Leiden clustering, 3D spatial graph formation, and graph feature extraction for risk stratification. We evaluated the results across four aspects: downstream prognostic performance, computational efficiency, preservation of phenotypic information and robustness to noise (**Fig. 4a**).

**Figure 4.**
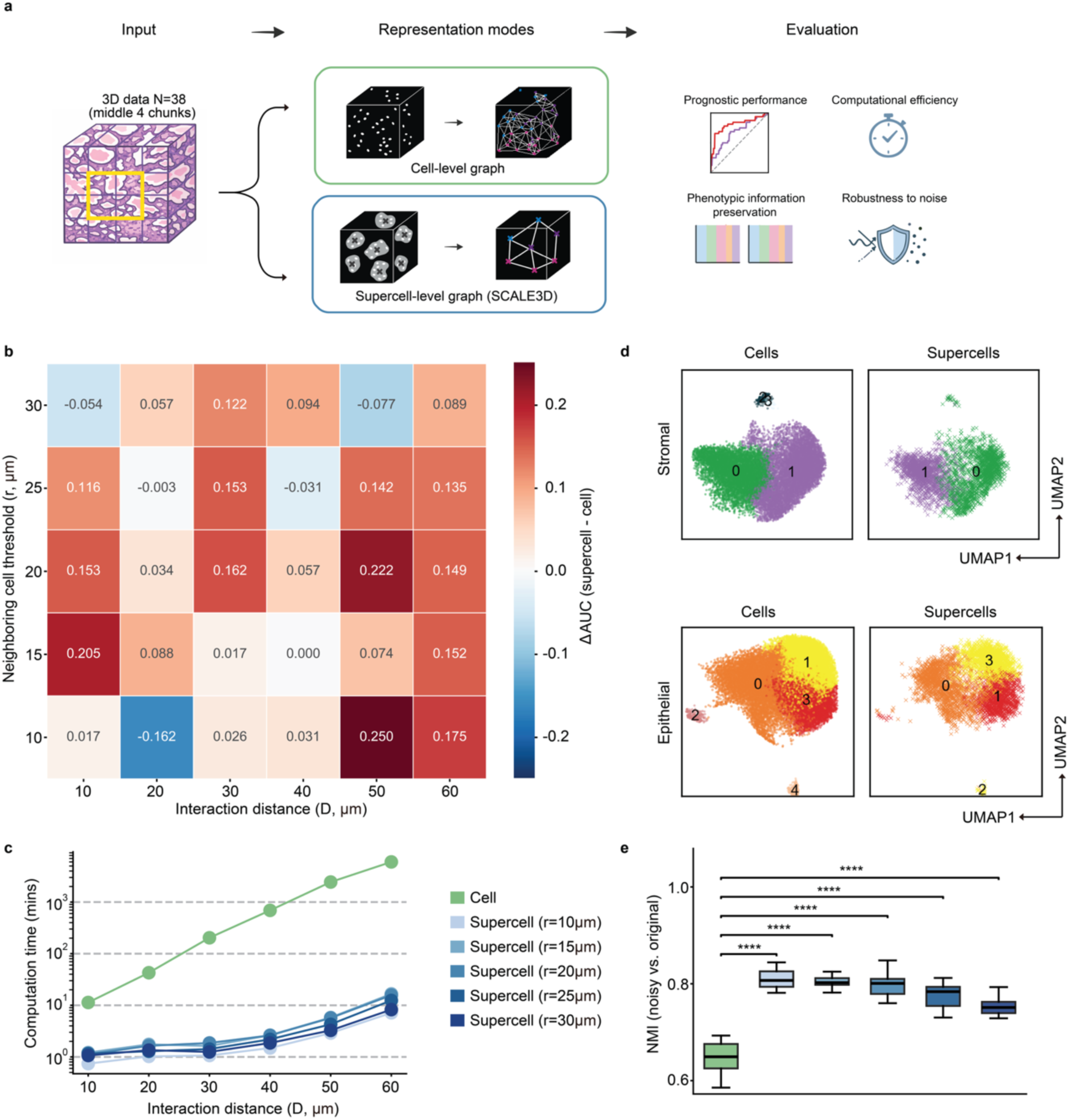
SCALE3D versus cell-level graph analysis. (a) Schematic overview of the comparison framework. A subset of 3D pathology data (N=38, each represented by 25% of the original tissue volume) was processed using either a conventional cell-level graph pipeline or the proposed supercell-based SCALE3D pipeline. Both representations underwent identical downstream steps, including morphology-based subtyping (Leiden clustering), 3D spatial graph formation, and graph feature extraction, followed by evaluation of predictive performance, computational efficiency, phenotypic preservation, and robustness to noise. (b) Heatmap showing the difference in predictive performance (ΔAUC: supercell−cell) for 5-year BCR across combinations of neighboring cell-distance thresholds (r, 10–30 µm) and interaction distance thresholds (D, 10–60 µm). Positive values (red) indicate improved performance of SCALE3D, observed across most parameter combinations. (c) Computational time for graph formation and graph feature extraction using cell-level and supercell representations (r, 10–30 µm) across interaction distances (D, 10–60 µm). SCALE3D substantially reduces computational cost across all configurations, with speedups increasing from approximately 10-fold to over 1000-fold as D increases. (d) Preservation of phenotypic structure. UMAP embeddings of cell-level (left) and supercell-level (right) morphometric features for stromal (top) and epithelial (bottom) compartments. For supercells, UMAP coordinates were computed as the mean of their constituent cells. Despite reduced granularity, supercells preserve the dominant clusters and overall feature space observed at the single-cell level. (e) Robustness to feature noise. Boxplots showing normalized mutual information (NMI) for Leiden clustering based on noisy versus original nuclear features (10 replicates, α = 0.1), for both cell-level (green) and supercell representations (blue) across r = 10–30 µm. Supercell representations consistently achieve higher NMI than cell-level representations, indicating greater stability to feature perturbations. Statistical significance is assessed using paired two-sided t-tests (****adjusted P < 0.0001)). Exact test statistics and P values are reported in Extended Data Table 5.

First, we evaluated predictive performance for 5-year BCR using LASSO regression under a LOOCV framework. Under the previously identified optimal configuration (r = 15 µm, D = 20 µm), SCALE3D features achieved a better performance over cell-level graph features (ΔAUC = 0.088; **Fig. 4b**). To ensure robustness, we evaluated ΔAUC across a range of r (10–30 µm) and D (10–60 µm). Overall, ΔAUC was positive across most parameter combinations, revealing a consistent trend toward comparable or higher performance with SCALE3D relative to cell-level analysis (**Fig. 4b**).

This pattern was most apparent at intermediate neighboring-cell thresholds (r=15–20 µm), where SCALE3D showed comparable or higher performance across all interaction distances (**Fig. 4b**). In contrast, smaller (r = 10 µm) and larger (r = 30 µm) thresholds showed more variable results, including occasional decreases in performance, suggesting that both overly fine and overly coarse representations may reduce predictive performance (**Fig. 4b**).

Meanwhile, we recorded the computation time for graph formation and graph feature extraction across all experiments. SCALE3D greatly reduced computational cost across all supercell formation parameters ( r = 10–30 µm). Notably, as the interaction distance increased (D = 10–60 µm), the speedup of the supercell representation increased from approximately 10-fold to over 1,000-fold compared to the cell-level approach (**Fig. 4c**).

Next, we evaluated whether supercell formation preserves morphology-defined phenotypic structure. Phenotypic preservation was quantified by comparing Leiden clustering of individual cells and supercells using normalized mutual information (NMI)³⁵, which measures how similar the clustering results are between the two representations. Across supercell formation parameters (r = 10–30 µm), NMI values consistently exceeded 0.65 (**Extended Data Table 4**), indicating moderate to strong agreement given the change in representation from cells to coarse-grained supercells. Complementary UMAP visualizations further support this observation. Supercells, projected to the mean UMAP coordinates of their constituent nuclei, occupy similar regions within the original cell-level feature manifold for both stromal and epithelial cells. Although cell-level Leiden clustering yields a larger number of morphology-based subtypes, reflecting finer granularity, the dominant subpopulation structure is preserved (**Fig. 4d**). Together, these results demonstrate that supercell formation retains the overarching morphology-defined phenotypic structure of the tissue microenvironment while significantly reducing data size.

In practice, morphology features extracted from pathology images are subject to variability arising from imperfect segmentation, staining heterogeneity, and imaging inconsistencies, which may affect downstream prediction tasks. To assess the robustness of supercell formation to perturbations in cell-level features, we added Gaussian noise with varying amplitudes (α = 0.05, 0.1, 0.2) to cell-level features across 10 replicates. Noisy supercells were then formed from the noisy cells, and Leiden clustering was performed independently for each representation. NMI metrics were calculated to compare the clustering results based on noisy versus original nuclear features (i.e. noisy supercells vs original supercells, noisy cells vs original cells). Across all noise amplitudes, supercell-level clustering consistently exhibits higher NMI than cell-level clustering (paired two-sided t-test, p < 0.001; **Fig. 4e; Extended Data Fig. 5; Extended Data Table 5**), indicating greater robustness to feature noise.

## Discussion

In this study, we introduce and validate SCALE3D, a data-efficient computational framework for modeling broad cellular interactions in large-scale 3D pathology datasets. The core idea is to use a biologically motivated coarse-grained strategy that groups morphologically similar neighboring nuclei into “supercells”. Preliminary clinical validation with prostate cancer specimens further demonstrates that cell-interaction features extracted from the SCALE3D pipeline provide meaningful predictive value for 5-year BCR outcomes and offer complementary prognostic information to established 3D glandular and nuclear morphological features.

Notably, an interaction distance of approximately 20 µm yields the best performance for BCR prediction. This finding is consistent with prior 2D spatial-omics and cell-graph studies suggesting that short-range cellular interactions play a critical role in shaping prostate cancer behavior^4,45^. In the final optimized model, the most discriminative features involve epithelial–epithelial and epithelial–stromal interaction patterns. In particular, patients who experience recurrence show greater variability in how epithelial cells are spatially organized, suggesting a more uneven and disordered tissue architecture. Differences are also observed in how epithelial cells interact with stromal cells, highlighting the potential importance of the surrounding stroma in disease progression.

Compared with standard cell-level graph analysis, SCALE3D reduces graph complexity while preserving overarching phenotypic and spatial interaction information. Importantly, it achieves comparable and, in some parameter settings, higher predictive performance relative to individual cell-level approaches. One possible explanation for this improved performance is that supercell-level representations reduce noise and redundancy in the feature space. When Gaussian noise is introduced at the cell level, our results show that supercell-level subtyping is significantly more stable than cell-level clustering. This suggests that supercell formation may act as a denoising mechanism, producing more robust representations and thereby reducing overfitting in small cohorts. In addition, coarse-grained representations may also reduce redundancy in cell-level graphs, emphasizing dominant spatial patterns while suppressing spurious interactions.

SCALE3D also provides substantial computational gains, particularly as the interaction distance between cells/supercells increases. At the cell level, computational cost grows rapidly—e.g. from minutes to thousands of minutes—as the interaction distance expands. In contrast, supercell-level representations scale much more efficiently, with computation times increasing only modestly (e.g. from minutes to tens of minutes). This improved scalability enables the exploration of a wider range of spatial interaction scales. This is important since the optimal interaction scale may vary across disease types, tissue architectures, and experimental settings.

Compared with immunolabeling and spatial transcriptomics, which are common ways to study cell interactions, our 3D pathology workflow offers several advantages. First, it enables scalable imaging of intact tissue volumes rather than thin 2D sections or shallow depths. This is important as our results, and prior studies, show that incorporating larger tissue volumes improves diagnostic performance by better capturing spatial heterogeneity and volumetric context^15,44,46,47^. In addition, this approach offers practical advantages in speed and cost: our fluorescent small-molecule analog of H&E staining yields rapid, uniform labeling of large tissues^14^, whereas immunolabeling relies on expensive antibodies with slow and often incomplete thick-tissue penetration. Finally, H&E-like contrast closely resembles conventional histopathology, facilitating interpretability and clinical acceptance.

Despite these strengths, several limitations exist and offer opportunities for further refinement. First, our current framework relies on nuclear morphology–based supercell subtyping, which may limit biological interpretability in the absence of molecular validation. Furthermore, nuclear morphology-based clustering may be sensitive to feature selection and algorithmic parameters. Future integration with multiplexed spatial transcriptomics or proteomics could refine cellular subtyping and enable mechanistic interpretation of interaction features. In addition, incorporating artificial intelligence approaches, such as deep representation learning or active learning frameworks in collaboration with pathologists, could further improve cell-type classification accuracy and robustness^48–50^. Second, 3D analysis is performed on subdivided tissue chunks due to computational constraints. Future work incorporating optimized graph partitioning could enhance whole-volume integration. Third, due to the relatively recent emergence of 3D pathology, the current study is based on a retrospective prostatectomy cohort from a single institution, which may limit generalizability across diverse patient populations and treatment settings.

In summary, we present a data-efficient computational pipeline for quantifying cellular interactions in large-scale 3D pathology datasets. This is achieved by introducing biologically informed supercell formation and scalable graph formation. These proof-of-concept findings suggest that 3D spatial graph analytics may complement existing 3D spatial biomarkers for improved risk stratification.

## Methods and Materials

### Tissue preparation and 3D imaging

Archived FFPE prostatectomy specimens from 76 patients with known clinical outcomes were obtained from the University of Pennsylvania. Patients were followed for at least 5 years after radical prostatectomy, with 5-year biochemical recurrence (BCR) status and time-to-BCR used as study endpoints. A genitourinary pathologist (P.L.) reviewed each case and selected a tumor-representative region, from which a single 3-mm-diameter punch biopsy (∼0.5-mm thickness) was extracted per case. Each punch biopsy was labeled with a small-molecule fluorescent analog of H&E consisting of TOPRO-3 Iodide (1:500 dilution) for nuclear staining and Alcoholic Eosin Y (1:100 dilution) for cytoplasmic staining, according to a previously published protocol^14^. Stained tissues were then dehydrated and cleared in ethyl cinnamate for imaging. 3D imaging was performed using an open-top light-sheet (OTLS) microscope. H&E-analog–stained prostate punches were imaged using 561-nm (eosin) and 638-nm (TO-PRO-3) excitation. Stitched and fused 3D volumes were generated using BigStitcher in ImageJ, followed by flat-field correction (stripe correction and depth adjustment) and intensity normalization using established post-processing pipelines^14^.

### Nuclear segmentation and morphological feature extraction

Nuclei were segmented using our previously established 3D image analysis pipelines^44^. Given the memory requirements necessary for 3D segmentation, fused 3D volumes were broken into 16 chunks for processing. Each chunk was 1024 × 1024 × 500 voxels in size, or approximately 540 × 540 × 270 µm. Prior to segmentation, each chunk underwent Laplacian-of-Gaussian sharpening for nuclear boundaries enhancement, 3D median filter using a spherical structuring element (r = 2 voxels) for denoising, and contrast-enhanced by adaptive histogram equalization (CLAHE) for local contrast variation correction. Preprocessed chunks were then segmented from using Cellpose with an estimated average nuclear diameter of 18 voxels and the pretrained denoise_nuclei model^34^. To prevent downstream quantification of nuclei that were artificially fragmented or cropped at chunk boundaries, segmentation masks were post-processed by removing border-touching objects. In addition, small spurious detections were excluded by filtering nuclei with volumes below the 1st percentile.

Quantitative nuclear morphological features including shape, intensity, texture and spatial context were extracted using a custom Python pipeline built on SciPy^51^ and scikit-image^52^. Shape features (e.g., nuclear volume, sphericity, and major/minor axis lengths) were computed directly from 3D nuclear segmentation masks. Cytoplasmic context was defined by dilating each nuclear mask with a spherical structuring element (radius = 10 voxels). Intensity-based features included first-order statistics (mean, standard deviation, minimum, and maximum) computed for both nuclear and cytoplasmic regions. Higher-order texture features were derived from gray-level co-occurrence matrices (GLCMs), computed on three orthogonal planes intersecting the nuclear centroid, with summary statistics (mean, standard deviation, minimum, maximum, and median across planes) aggregated per nucleus. In addition, nuclear intensity entropy was calculated from a normalized 32-bin intensity histogram. Spatial contextual features included local cellular crowdedness, quantified by constructing a k-nearest neighbor graph (k = 15) using a KD-tree and computing the per-nucleus mean and standard deviation of neighbor distances. Nuclear centroid coordinates were recorded for downstream analyses. Previously generated glandular masks^15^ were used to classify segmented nuclei as belonging to epithelial or stromal compartments based on the centroid location. A full feature list is provided in **Extended Data Table 1**.

### Supercell formation

To reduce data complexity while preserving key cell–cell interaction patterns, individual nuclei were grouped into supercells based on spatial proximity and morphological similarity. Nuclei were first stratified by tissue compartment (epithelial or stromal) based on precomputed glandular mask^43^, and supercell formation was performed independently within each region. Each nucleus was represented by 52 morphometric features extracted in the previous step. Before computing feature similarity, each feature dimension was standardized using the mean and standard deviation calculated across all datasets. Supercells were then generated using an iterative region-growing procedure. Starting from a randomly selected nucleus, neighboring nuclei were evaluated as candidate members of the growing supercell. A candidate nucleus was added only if it satisfied two criteria: (i) its spatial centroid was within radius r of the original seed nucleus, and (ii) the cosine similarity between its standardized feature vector and the current mean feature vector of the growing supercell was ≥0.7. After each accepted nucleus, the supercell mean feature vector was updated. Supercell growth continued until no additional candidate nuclei satisfied the inclusion criteria or until the radius-specific maximum supercell size was reached. This maximum size was defined separately for each neighboring-cell threshold r, based on the mean number of nuclei identified within that radius across all datasets. Candidate supercells containing fewer than three nuclei are discarded. This process is repeated until every nucleus in every 3D chunk was either assigned to a supercell or considered during the algorithm. All 3D chunks were processed in this manner. For each resulting supercell, features were averaged from its constituent nuclei and the centroids were computed as the mean spatial coordinates.

### Supercell subtyping

Supercell subtypes were then identified by applying Leiden clustering to the pooled supercell-level morphological feature vectors across chunks and specimens. All morphological features were standardized and reduced in dimensionality using principal component analysis (PCA). To correct for specimen-specific batch effects, the PCA embeddings were harmonized using Harmony, yielding a batch-corrected latent space. A k-nearest neighbor graph was constructed in the Harmony-corrected space using 20 principal components and 50 neighbors. For visualization and unsupervised subtype discovery, UMAP embeddings were computed from the Harmony neighbor graph, and Leiden community detection was applied at a resolution of 0.1. This pipeline was implemented using a GPU-accelerated single-cell analysis workflow based on Scanpy^53^ and rapids-singlecell^54^.

### Graph formation and graph feature extraction

To characterize spatial supercell interactions, a spatial neighbor graph was constructed for each data chunk using squidpy.gr.spatial_neighbors^55^, based on Euclidean distances between supercell centroids within a threshold (D). This resulted in a spatial graph in which nodes represent supercells and edges represent spatial proximity within the specified radius. From each chunks, supercell interaction features were computed using Squidpy^55^, including (i) interaction frequency matrices quantifying the number of spatial contacts between each pair of supercell subtypes, (ii) neighborhood enrichment z-scores measuring over- or under-representation of subtype interactions relative to random expectation, and (iii) supercell subtype-specific centrality measures the relative importance of each subtype within the graph. In addition, global graph statistics (e.g. node count, average node degree, and graph density) were also computed. A full graph feature list is provided in **Extended Data Table 2**. To obtain a specimen-level representation, a total of 320 features from the 16 chunks within each specimen were summarized using first-order statistics (mean, minimum, maximum, standard deviation, and median) across chunks.

### Prognostic analysis

Patients who developed biochemical recurrence (BCR) within 5 years after radical prostatectomy (RP) were labeled as BCR, while those with ≥5 years of follow-up without recurrence were classified as non-BCR. BCR was defined as a rise in serum prostate-specific antigen (PSA) to ≥0.2 ng/mL at least 8 weeks after RP.

We evaluated prognostic performance using two complementary approaches: (i) binary 5-year BCR prediction with LASSO logistic regression and (ii) time-to-event modeling with ridge-penalized Cox proportional hazards regression. Both analyses were conducted under a leave-one-out cross-validation (LOOCV) framework. Within each fold, preprocessing was performed exclusively on the training set to avoid data leakage. Features were standardized, highly correlated variables (Pearson r ≥ 0.9) were removed, and minimum redundancy–maximum relevance (mRMR) was used to select the top 20 features. The resulting scaling parameters and feature subset were then applied to the held-out sample.

For binary prediction, the LASSO regularization parameter (C) was tuned via inner 4-fold cross-validation on the training set to maximize AUC. The model was refit on the full training fold to generate out-of-fold (OOF) probabilities. Aggregated OOF predictions were used to compute AUC with 95% confidence intervals estimated by 1,000 bootstrap resamples. To assess overfitting, permutation testing (500 iterations) was performed by repeating the full pipeline with shuffled labels; the empirical p-value was defined as the fraction of permuted AUCs exceeding the observed AUC.

For survival analysis, ridge-penalized Cox models were trained within the same LOOCV framework, with penalization tuned by inner 4-fold cross-validation. OOF risk scores were used to stratify patients into high- and low-risk groups based on the median training-set score. Performance was evaluated using Kaplan–Meier analysis, log-rank test, hazard ratio (HR) with 95% confidence interval, and concordance index (C-index).

To assess the added prognostic value of SCALE3D-derived features beyond glandular and nuclear features, likelihood ratio tests were performed comparing Cox models fit to OOF risk scores. The reduced model included baseline features, while the full model additionally incorporated SCALE3D-based scores. Model fit was compared using the difference in Cox partial log-likelihoods, calculated as 2(*logL_full_* – *logL_reduced_*), and evaluated against a chi-square distribution with 1 degree of freedom. Univariate comparisons between BCR and non-BCR groups were conducted using the Mann–Whitney U test. All statistical tests were two-sided. For analyses involving multiple comparisons, P values were adjusted using the Benjamini–Hochberg false discovery rate procedure, and adjusted P values < 0.05 were considered statistically significant.

## Ethics Approval and Consent to Participate

Radical prostatectomy specimens used in this study were obtained from the University of Pennsylvania under Institutional Review Board (IRB) approval (#830205).

## Data availability

The image data are currently in the process of being deposited in The Cancer Imaging Archive (TCIA). The code used in this study is available at https://github.com/yzhao07/SCALE3D.

## Acknowledgements

This research was primarily supported by the National Cancer Institute (NCI) under R01CA268207 (Liu and Madabhushi), the National Institute of Diabetes and Digestive and Kidney Diseases (NIDDK) under R01DK138948 (Liu, Mahmood, and Grady), and the National Institute of Biomedical Imaging and Bioengineering (NIBIB) under R01EB031002 (Liu). NIH support for WM Grady includes U01CA152756, R01CA220004, U2CCA271902, U54CA163060, and U01CA182940. Funding for WM Grady is also provided by the Prevent Cancer Foundation, Cottrell Family Fund, Evergreen Fund, and Listwin Foundation to WMG. These studies are also supported by GiCaRes from UW Departments of Medicine (Gastroenterology Division) and Lab Medicine & Pathology. Additional support was provided by the Department of Defense (DoD) Prostate Cancer Research Program (PCRP) under W81XWH-20-1-0851 (Madabhushi and Liu), the Pacific Northwest Prostate Cancer SPORE P50CA97186 (True), and the Canary Foundation. The content is solely the responsibility of the authors and does not necessarily represent the official views of the National Institutes of Health, the U.S. Department of Veterans Affairs, the Department of Defense, or the United States Government.

