## Supplemental information for "Scalable 3D cell-interaction analysis via supercell graphs for prostate cancer risk stratification"

[1]*Department of Biomedical Informatics and Medical Education, University of Washington, Seattle, WA, USA*
[2]*Department of Pathology, Stanford University, Stanford, CA, USA*
[3]*Department of Mechanical Engineering, University of Washington, Seattle, WA, USA*
[4]*Department of Medicine, Section of Hematology/Oncology, University of Chicago, Chicago, IL, USA*
[5]*Department of Pathology, Mass General Brigham, Harvard Medical School, Boston, MA, USA*
[6]*Cancer Program, Broad Institute of Harvard and MIT, Cambridge, MA, USA*
[7]*Data Science Program, Dana-Farber Cancer Institute, Boston, MA, USA*
[8]Harvard-MIT Division of Health Sciences and Technology, Massachusetts Institute of Technology, Cambridge, MA, USA
[9]*Department of Translational Molecular Pathology, The University of Texas MD Anderson Cancer Center, Houston, TX, USA*
[10]*Institute for Data Science in Oncology, The University of Texas MD Anderson Cancer Center, Houston, TX, USA*
[11]*Pathology and Laboratory Medicine, University of Pennsylvania, Philadelphia, PA, USA*
[12]*Stanford Medical Center, Stanford, CA, USA*
[13]*Sunnybrook Health Sciences Centre, University of Toronto, Toronto, ON, Canada*
[14]*UC Berkeley-UCSF Graduate Program in Bioengineering, University of California, Berkeley, CA, USA*
[15]*Wallace H. Coulter Department of Biomedical Engineering, Emory University & Georgia Institute of Technology, Atlanta, GA, USA*
[16]*Harvard Data Science Initiative, Harvard University, Cambridge, MA, USA*
[17]*Department of Laboratory Medicine & Pathology, University of Washington, Seattle, WA, USA*
[18]*Department of Bioengineering, Stanford University, Stanford, CA, USA*
*

### Extended Data

#### Figures

Data-scale expansion and nuclei segmentation workflow

An image atlas of randomly selected 2D cross-sections from each specimen

Additional examples of the morphometric characterization shown in Fig. 3c

Differential epithelial–epithelial and epithelial–stromal interaction features between 5-year BCR and non-BCR groups.

Robustness to feature noise

#### Tables

A list of 3D nuclear morphological features

A list of 3D supercell graphs features

Statistical comparison of top 10 highly stable features between non-BCR and BCR groups

NMI across a range of supercell formation parameters (r=10-30 μm)

Statistical comparison of supercell- and cell-level NMI stability

#### Videos

Video abstract summarizing the SCALE3D study workflow and major findings.

Three-dimensional visualization of the SCALE3D workflow.

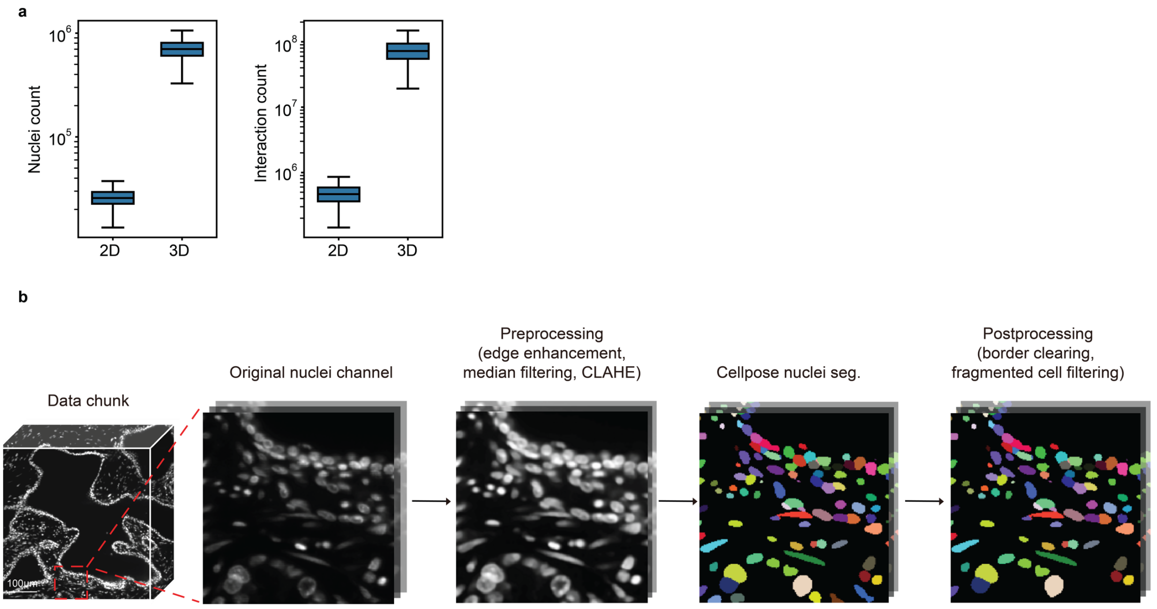


**Extended Data Figure 1.** Data-scale expansion and nuclei segmentation workflow. (a) Comparison of data scale between 2D and 3D pathology. Boxplots show the distribution of cell counts (left) and potential cell–cell interaction counts (right, radius graph distance = 40 μm) on a log scale. Transitioning from 2D sections to 3D volumetric imaging results in an approximate order-of-magnitude increase in the number of cells and a two-order-of-magnitude increase in potential cell–cell interactions, highlighting the computational complexity in 3D analysis. (b) Representative 2D cross-sectional views of the nuclei segmentation pipeline. The nuclei channel is used as input for segmentation. Preprocessing steps—including edge enhancement, median filtering, and contrast-limited adaptive histogram equalization (CLAHE)—are applied to improve signal quality. Nuclei are subsequently segmented using Cellpose to generate instance-level masks. Postprocessing procedures, including border clearing and removal of fragmented objects, are performed to further refine segmentation results.

AFM008

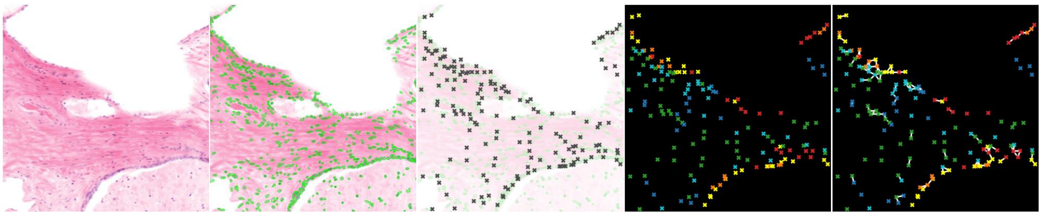

EAM003

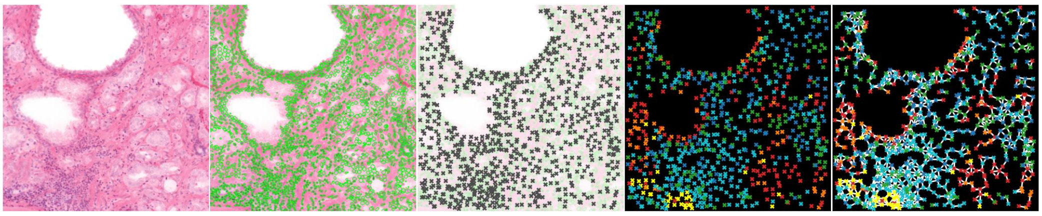

EAM004

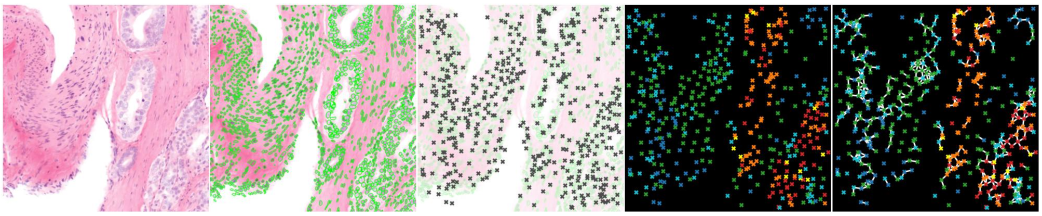

EAM012

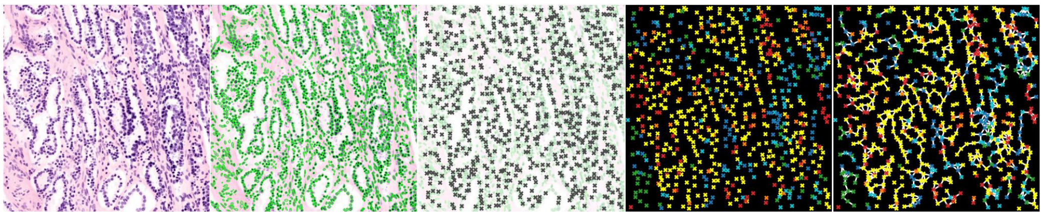

EAM017

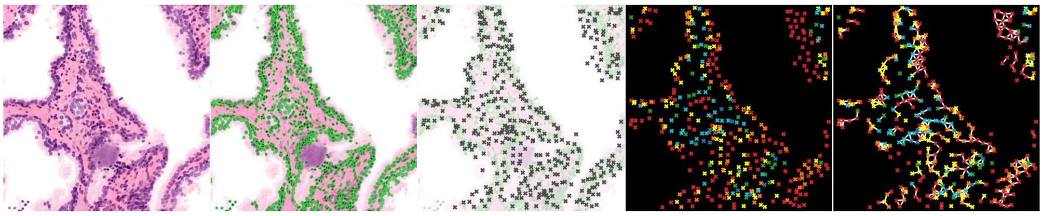

EAM018

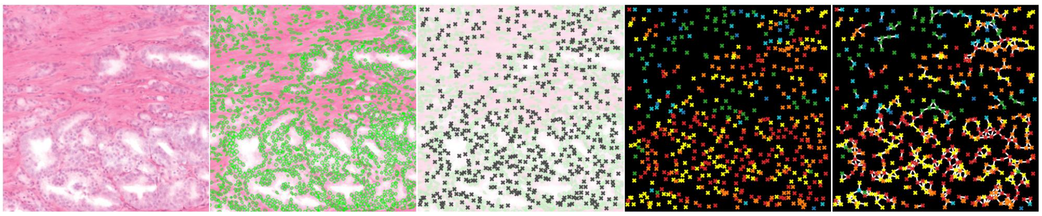

AFM001

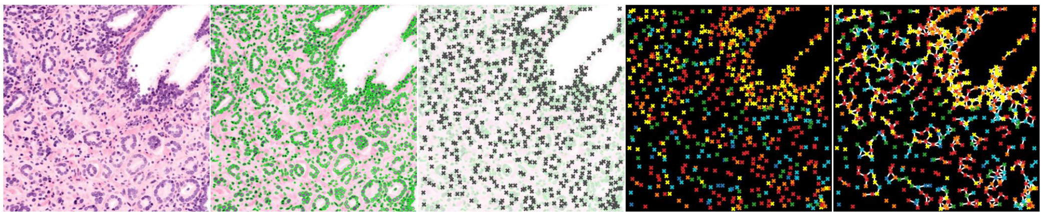

AFM021

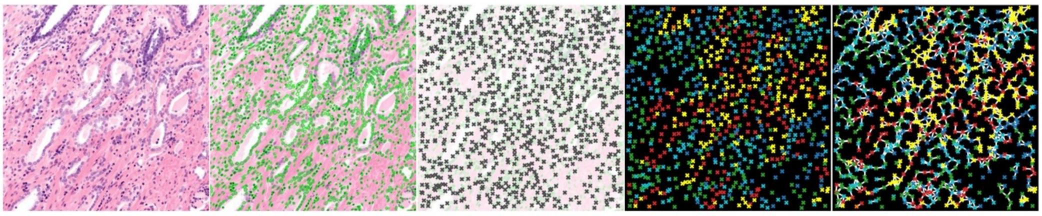

EAM020

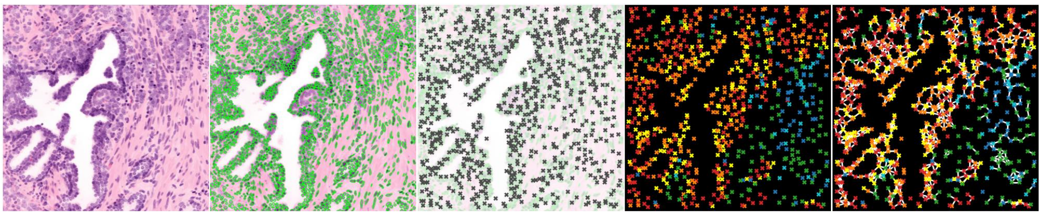

EAM021

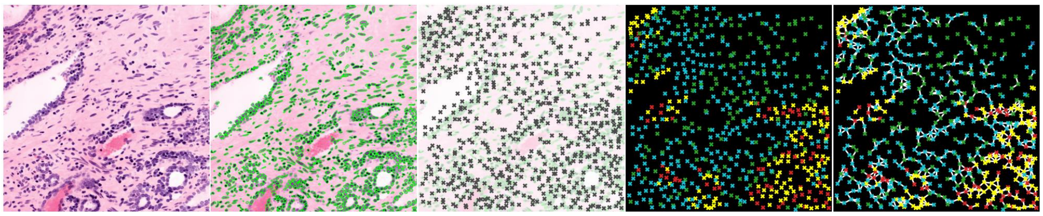

EAM019

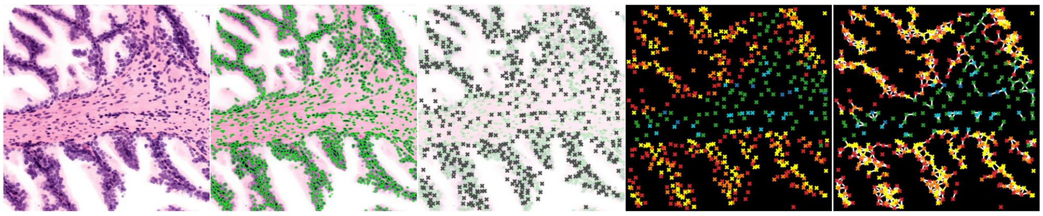

EAM022

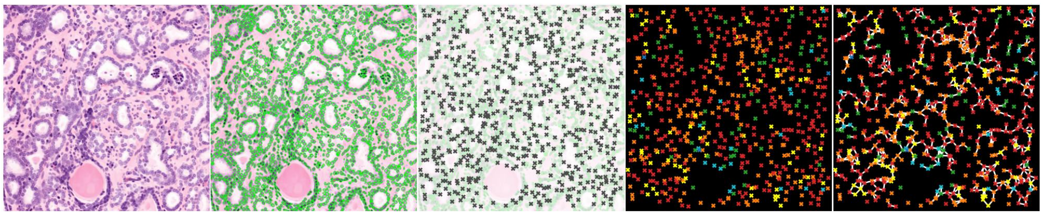

EAM033

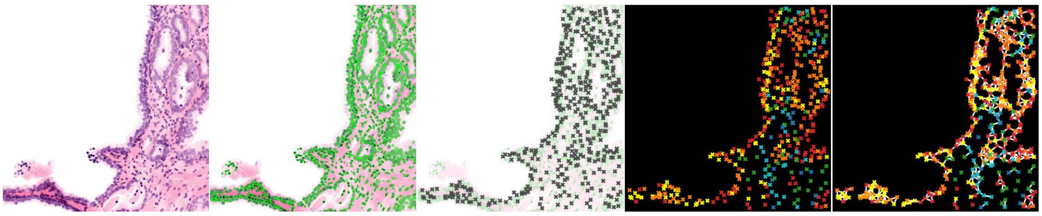

AFM025

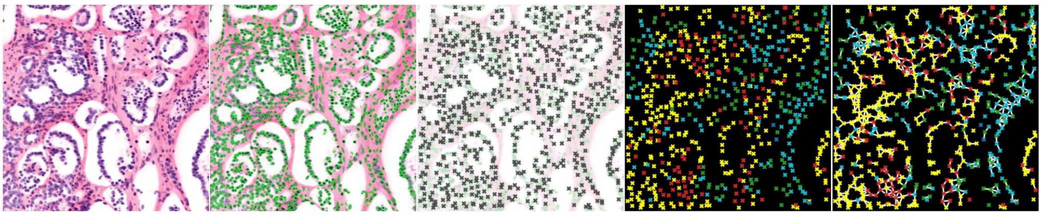

EAM035

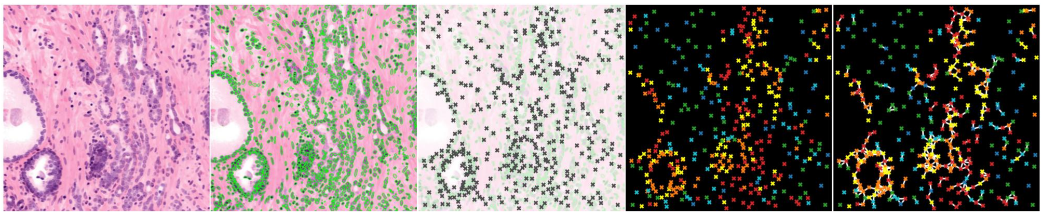

AFM030

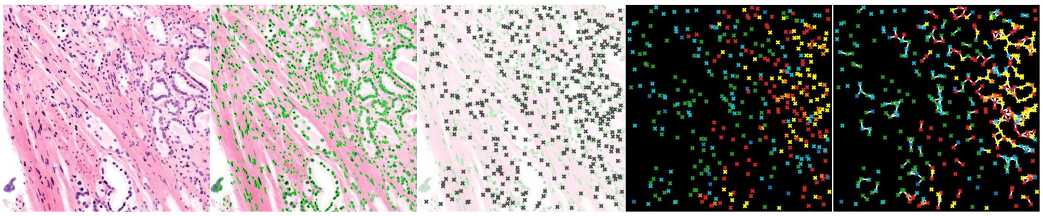

EAM037

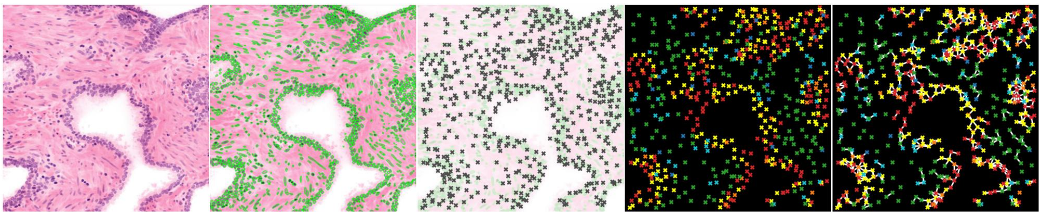

AFM054

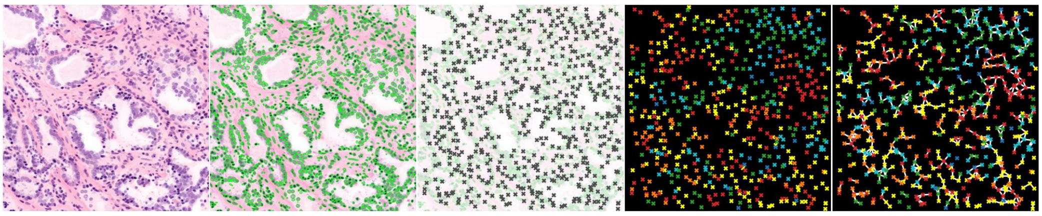

AFM059

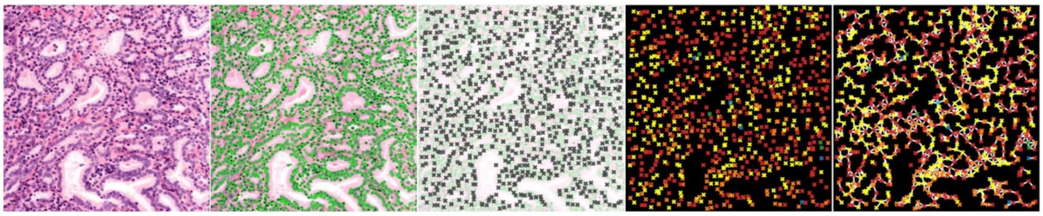

AFM064

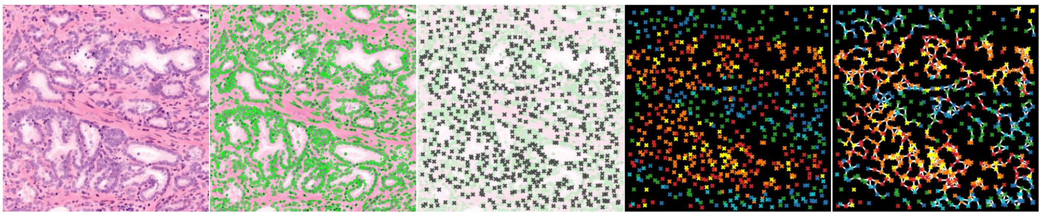

AFM033

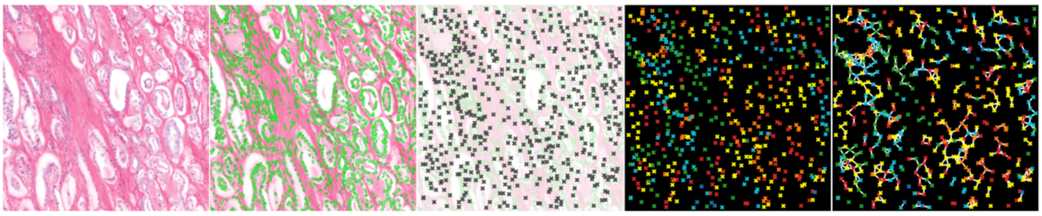

AFM039

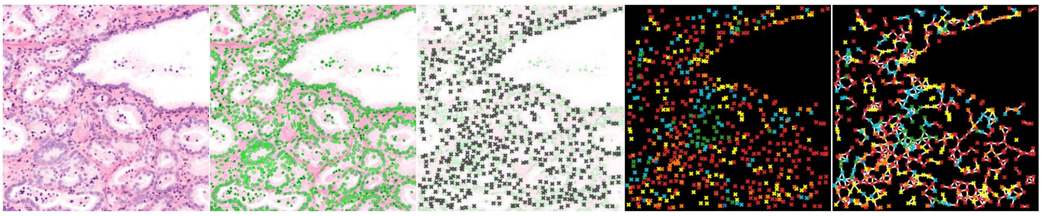

AFM042

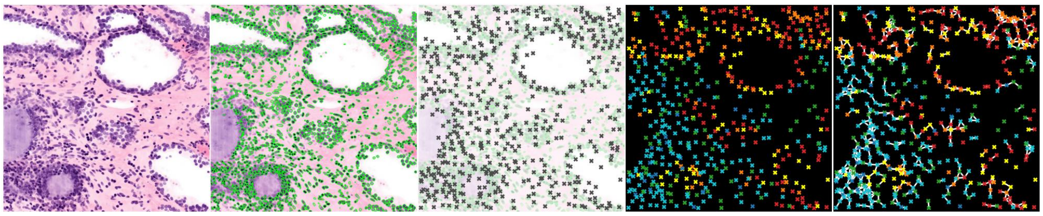

AFM046

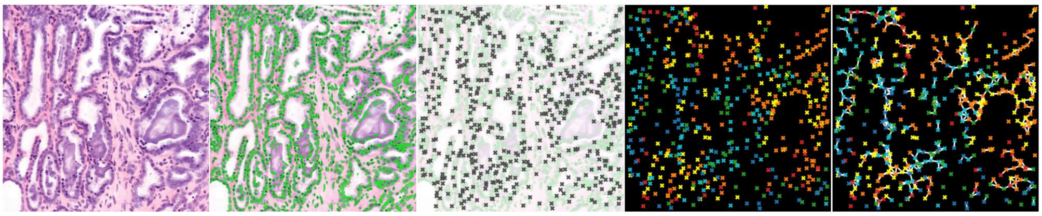

AFM047

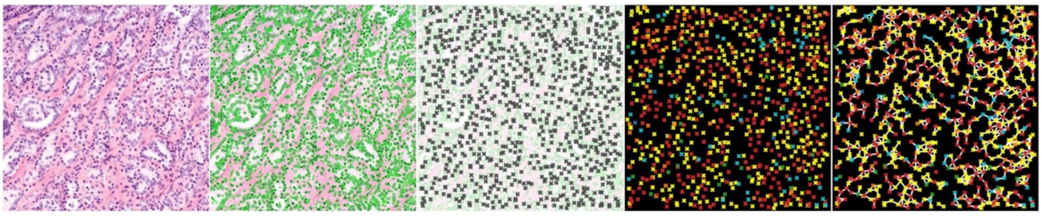

AFM050

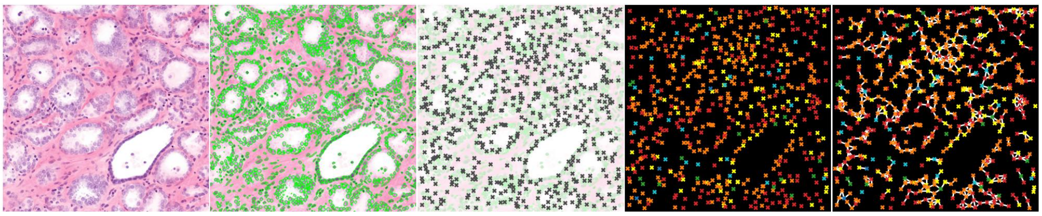

AFM051

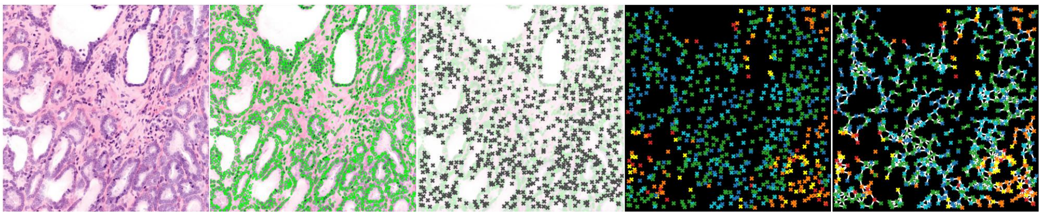

AFM052

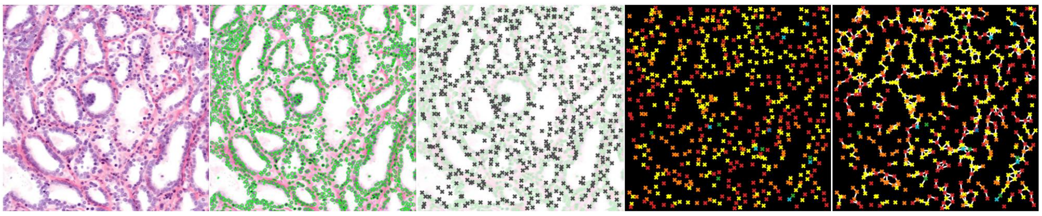

AFM053

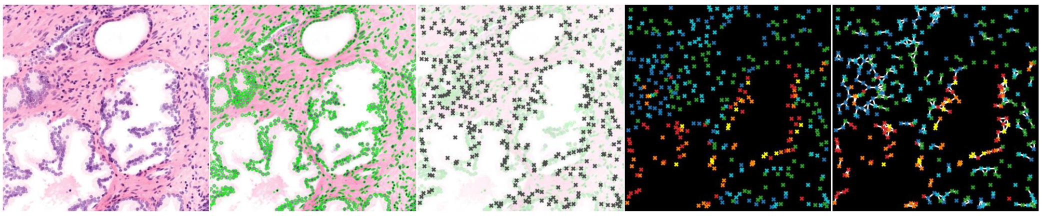

AFM066

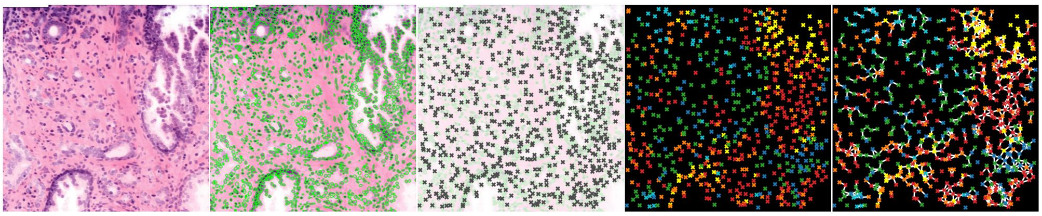

EAM049

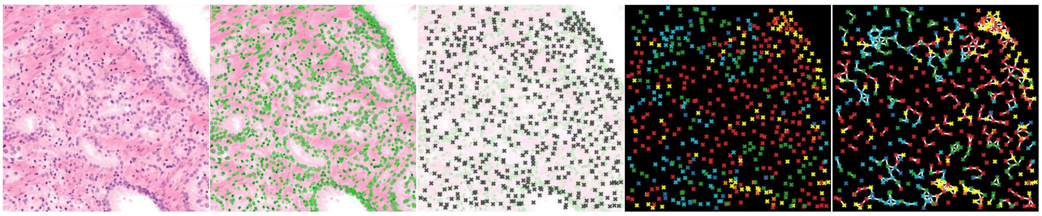

EAM052

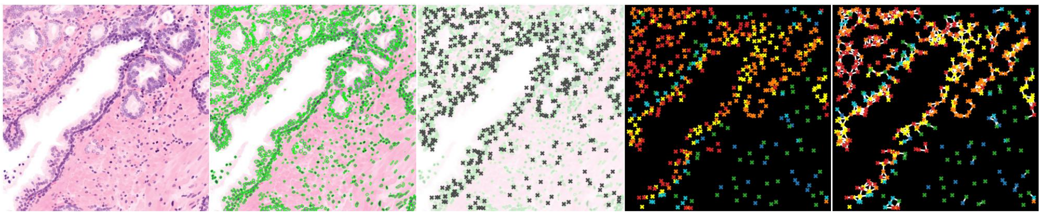

EAM053

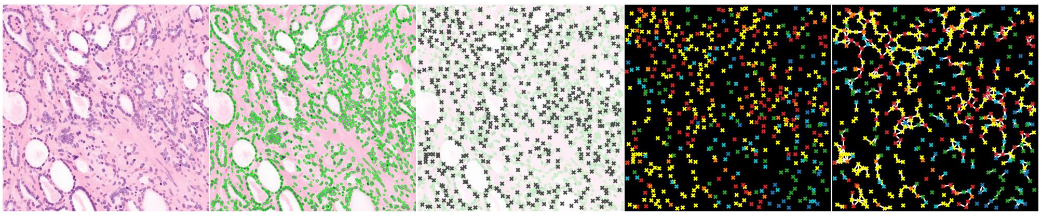

AFM071

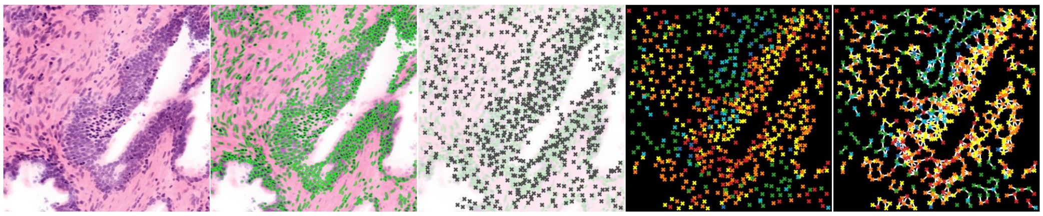

AFM072

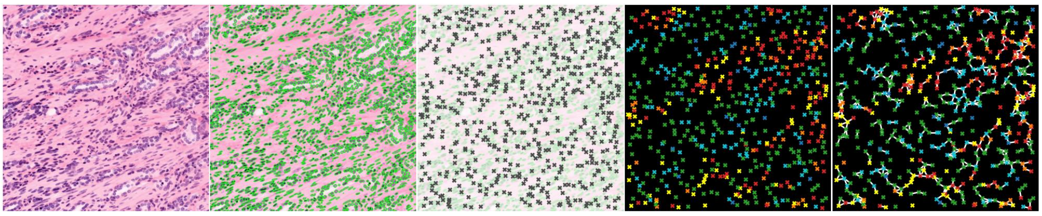

AFM073

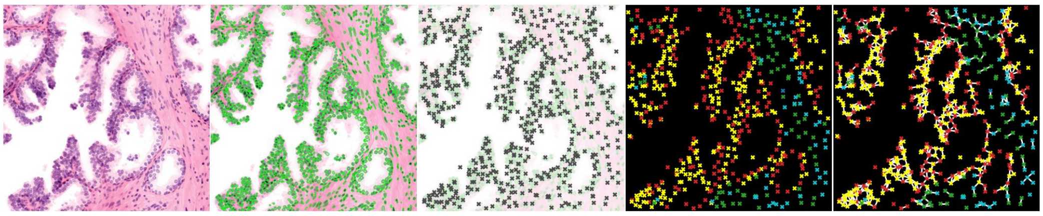
AFM074

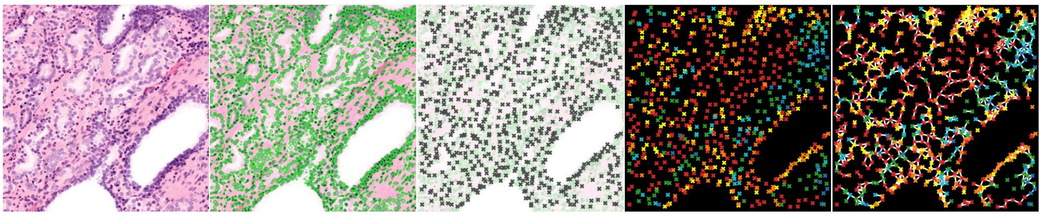
AFM076

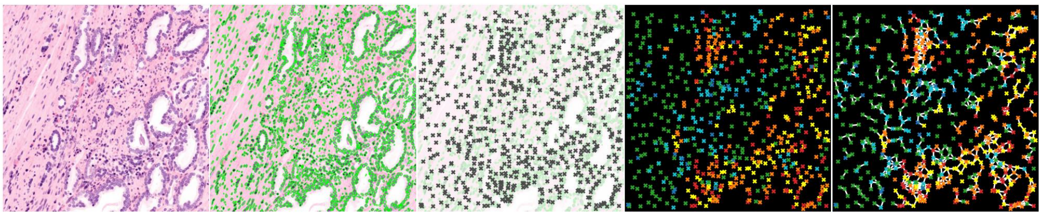
AFM077

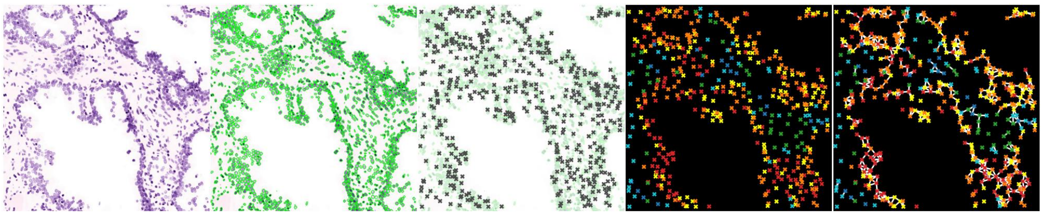
AFM078

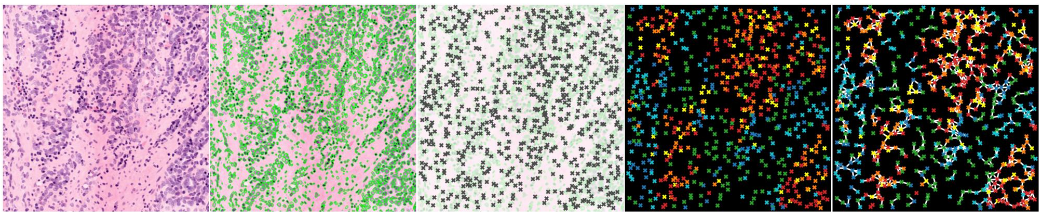
AFM080

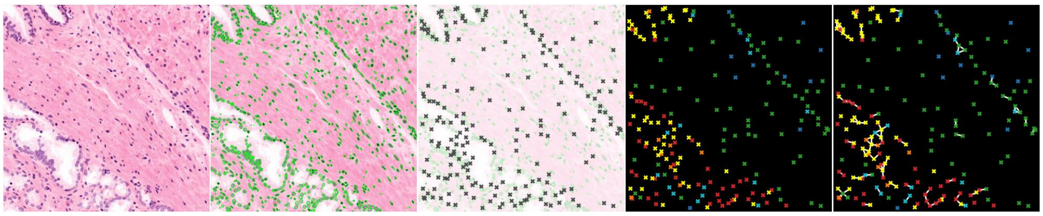
AFM081

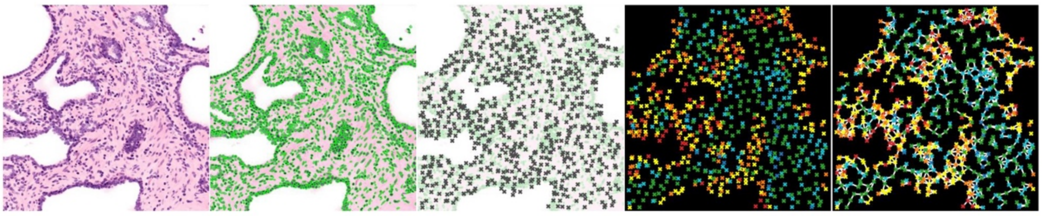
AFM082

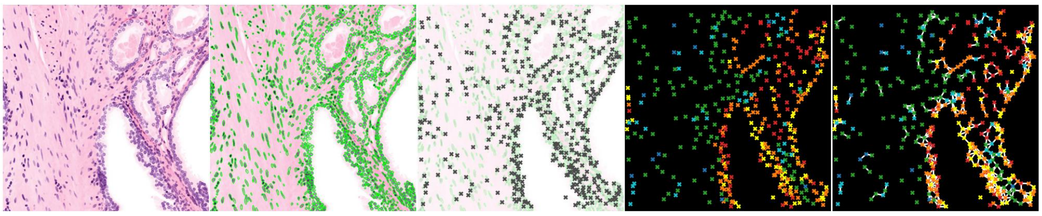

EAM056

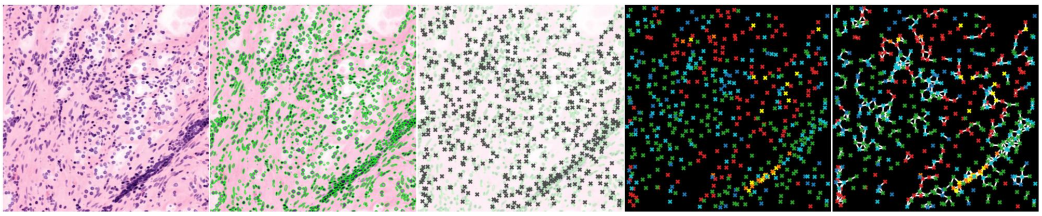

AFM068

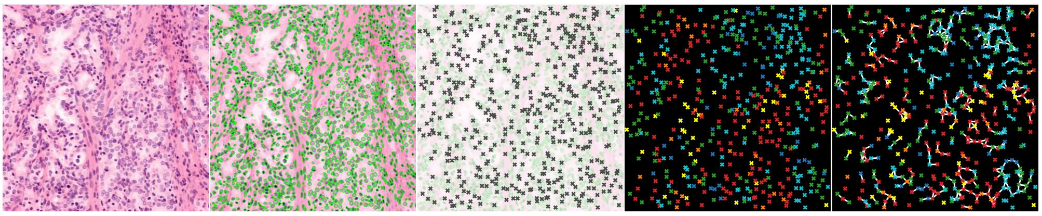

AFM083

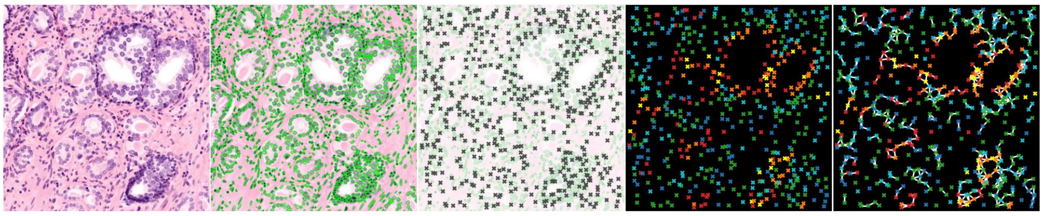

EAM057

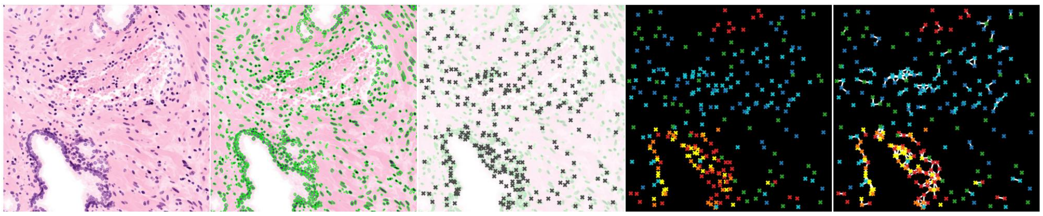

EAM060

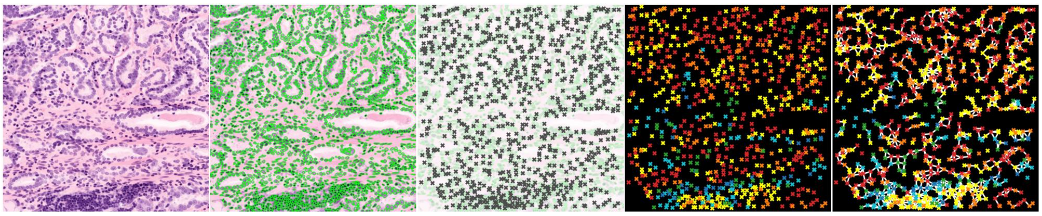

AFM086

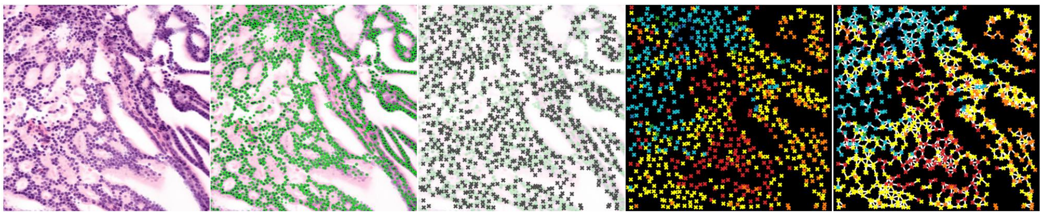

AFM069

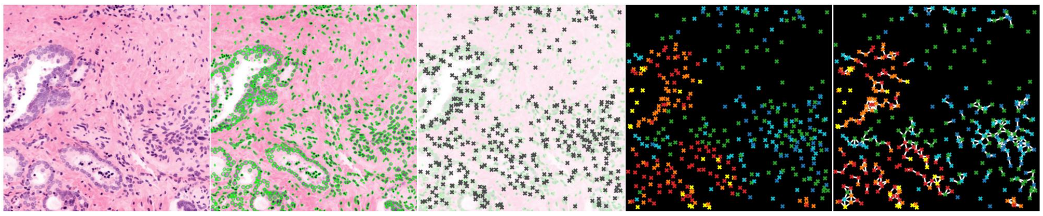
EAM041

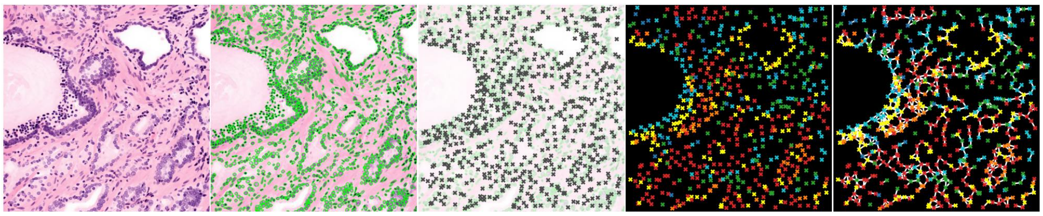
EAM043

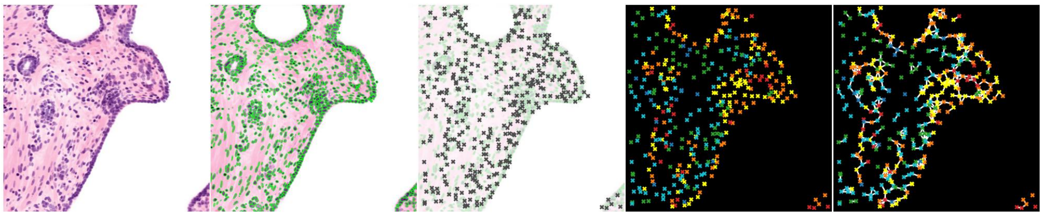
EAM044

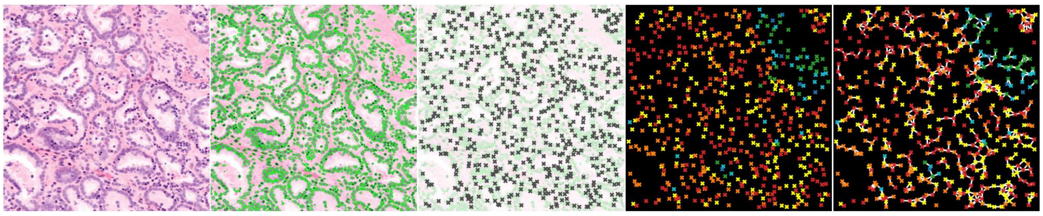
EAM046

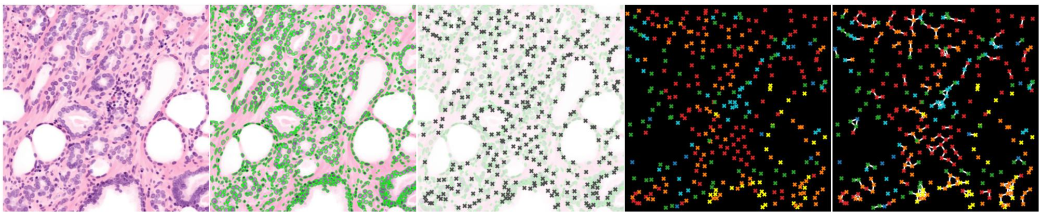
AFM101

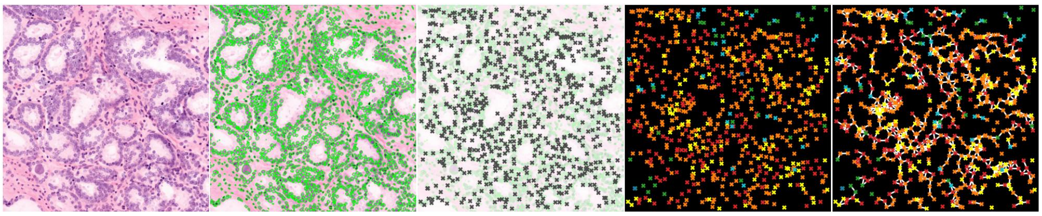
EAM078

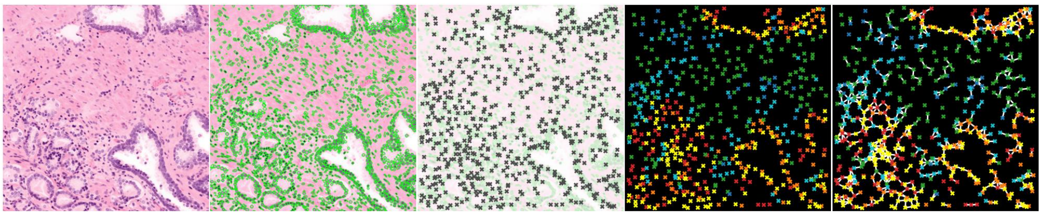
EAM079

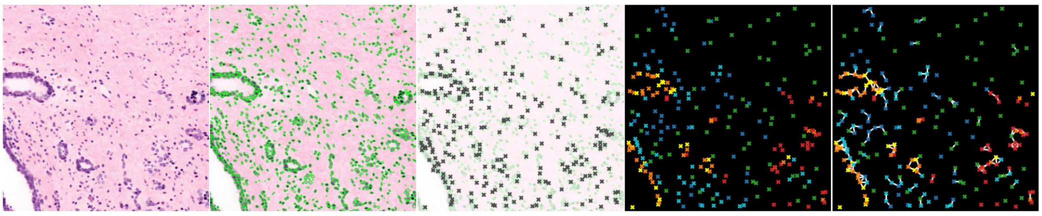

EAM084

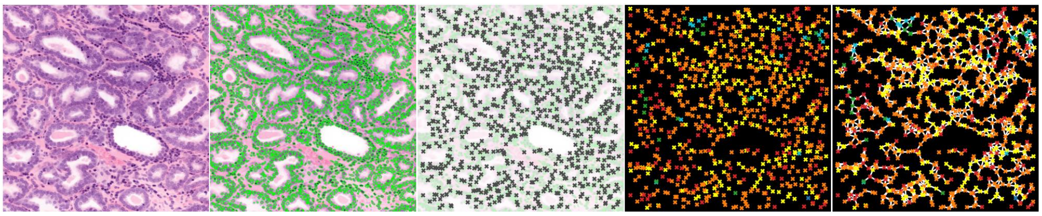

EAM085

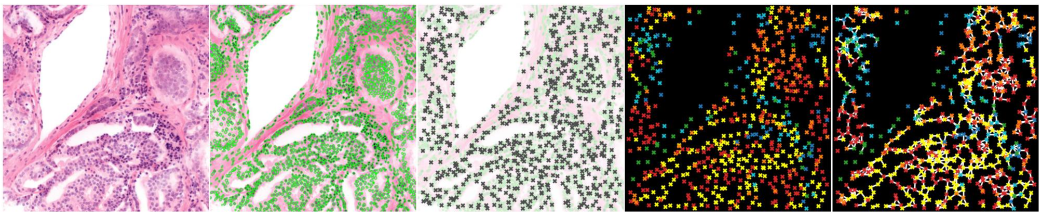

EAM087

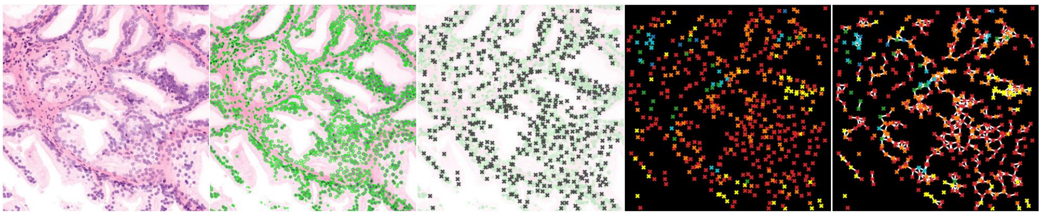

EAM088

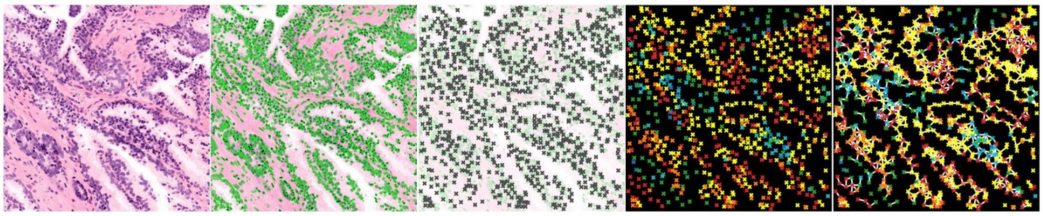

EAM089

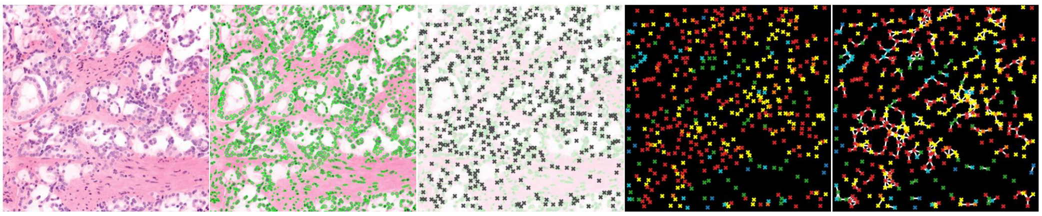

EAM098

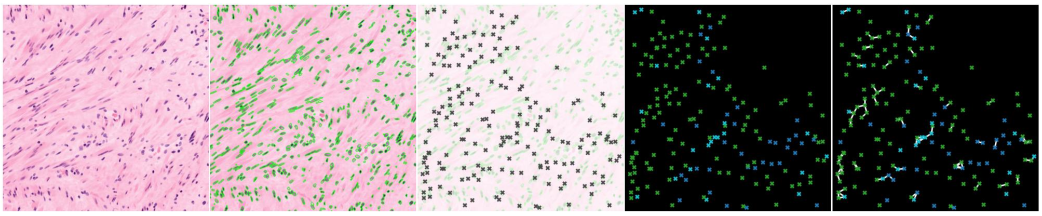

EAM062

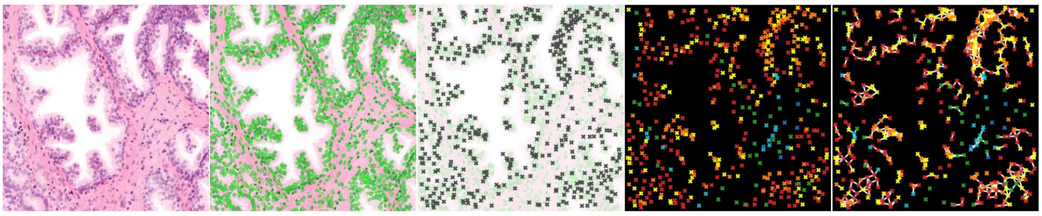

EAM063

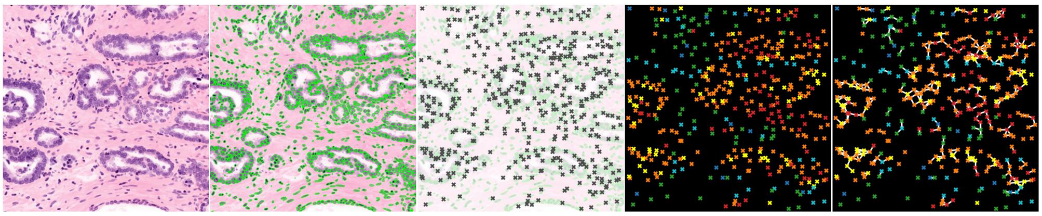

EAM069

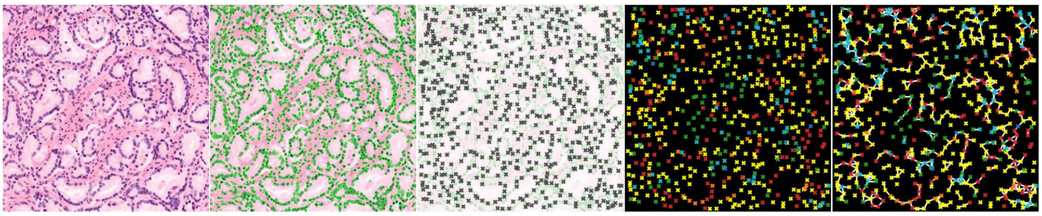

EAM071

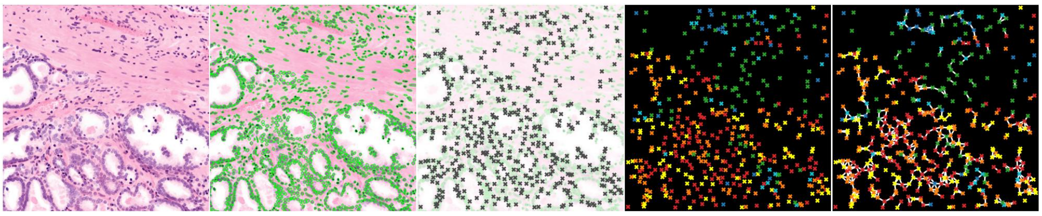

EAM076

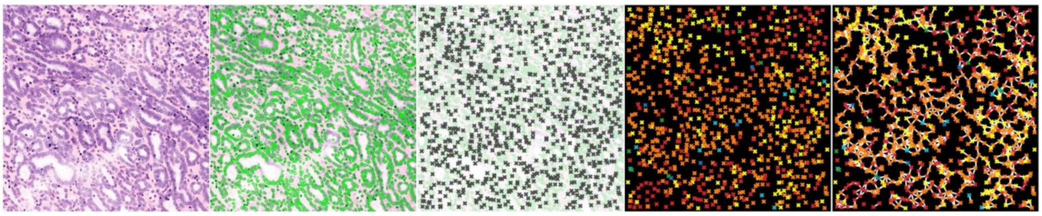

AFM097

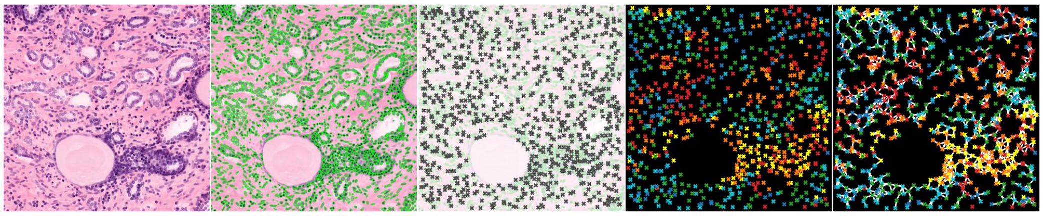

AFM099

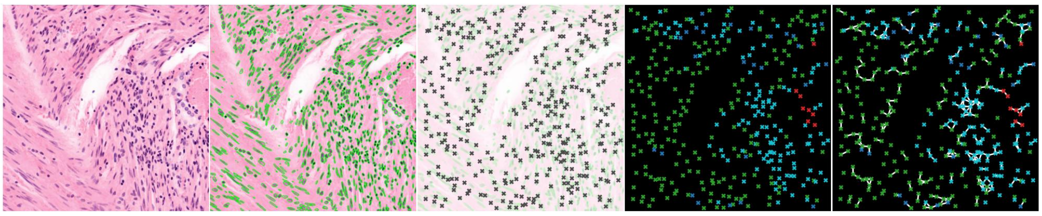

AFM012

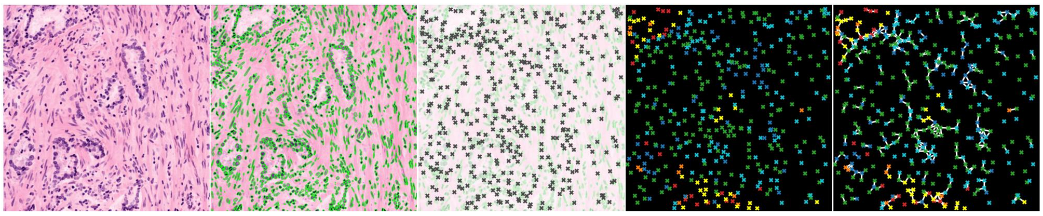

AFM014

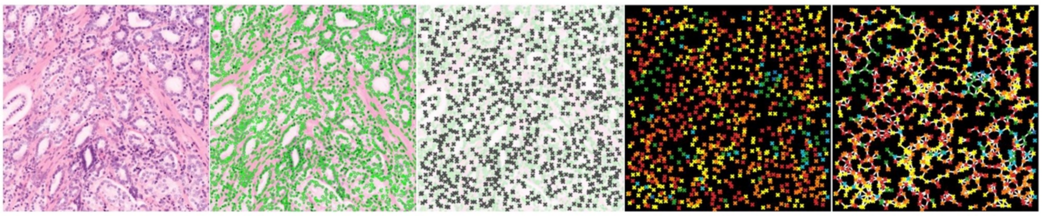

EAM002

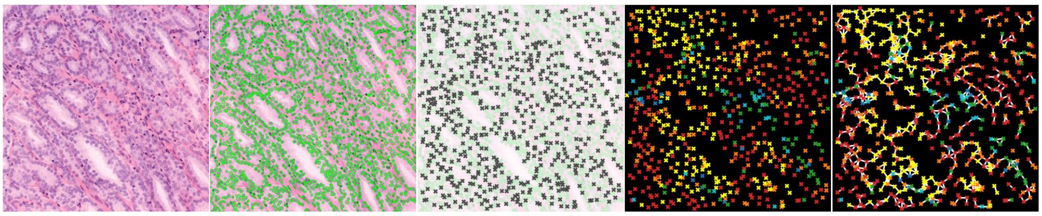

AFM023

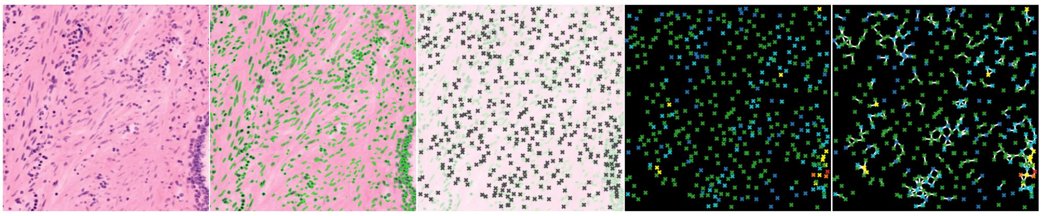

AFM024

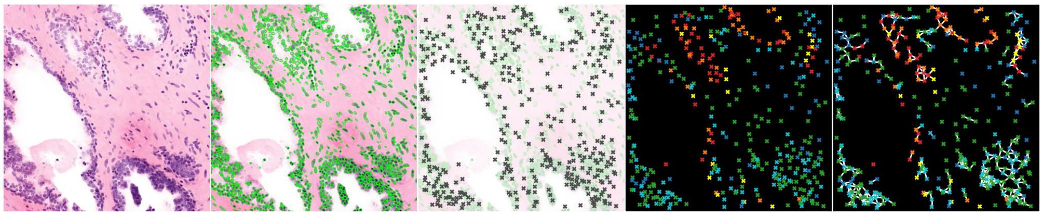

EAM009

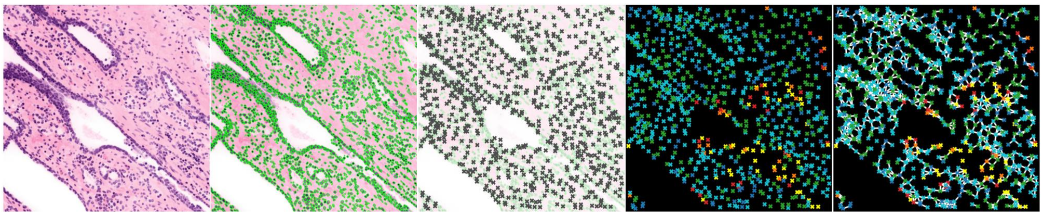

**Extended Data Figure 2.** An image atlas of randomly selected 2D cross-sections from each specimen, illustrating the chunk-level analysis pipeline, including H&E-analog images, nuclear segmentation, supercell formation, supercell subtyping, and supercell graph formation.

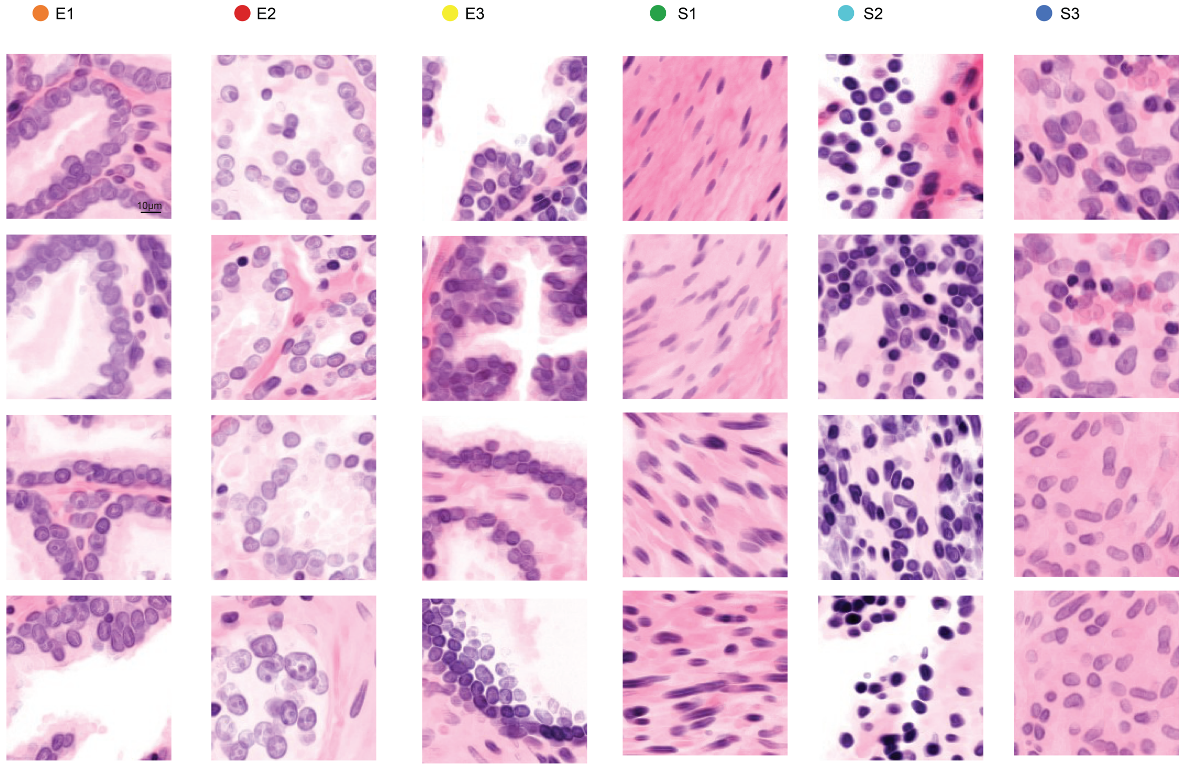


**Extended Data Figure 3.** Additional examples of the morphometric characterization shown in Fig. 3c for epithelial subtypes E1–E3 (orange, red, yellow) and stromal subtypes S1–S3 (green, cyan, blue).

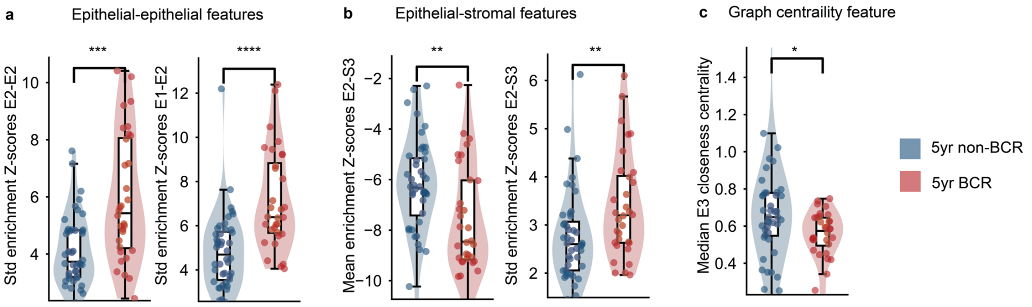


**Extended Data Figure 4.** Differential epithelial–epithelial and epithelial–stromal interaction features between 5-year BCR and non-BCR groups. (a) Epithelial–epithelial interaction variability, quantified by the standard deviation of enrichment Z-scores, is significantly higher in the BCR group for both E2-E2 and E2-E1 interactions, indicating increased heterogeneity in epithelial interaction patterns. (b) Epithelial–stromal interaction features also differ between groups: the mean enrichment Z-score for E2-S3 interactions is higher in the non-BCR group, whereas the variability of the same interaction is elevated in the BCR group. (c) The median E3 closeness centrality is significantly higher in the non-BCR group. Each point represents an individual specimen. Statistical significance was assessed using two-sided Mann–Whitney U tests. Significance is indicated based on Benjamini–Hochberg FDR-adjusted P values: *adjusted $P < 0.05$; **adjusted $P < 0.01$; ***adjusted $P < 0.001$; ****adjusted $P < 0.0001$. Exact P values are reported in Extended Data Table 3.

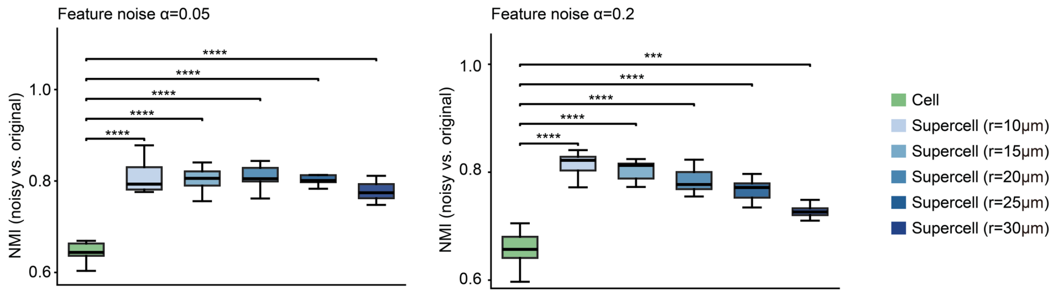


**Extended Data Figure 5.** Robustness to feature noise. Boxplots showing normalized mutual information (NMI) for Leiden clustering based on noisy versus original nuclear features (10 replicates; $\alpha = 0.05, 0.2$) for both cell-level and supercell representations across neighboring-cell thresholds r = 10–30 μm. Supercell representations consistently achieve higher NMI than cell-level representations, indicating greater stability to feature perturbations. Statistical significance was assessed using two-sided paired t-tests. Significance is indicated based on Benjamini–Hochberg FDR-adjusted P values: ***adjusted $P < 0.001$; ****adjusted $P < 0.0001$. Exact test statistics and P values are reported in Extended Data Table 5.

**Extended Data Table 1.** A list of 3D nuclear morphological features.

| Shape (11) | Intensity (8) | Texture (31) | Spatial information (2) |
|---|---|---|---|
| Volume, convex volume, Equivalent diameter, Extent, Major axis length, Minor axis length, Solidity, Surface area, Aspect ratio, Roughness, Sphericity | Nucleus intensity (std, mean, min, max), Cytoplasm intensity (std, mean, min, max), | GLCM_contrast (5), GLCM_dissimilarity (5), GLCM_homogeneity (5), GLCM_energy (5), GLCM_correlation (5), GLCM_ASM (5), Entropy | Crowdedness (std, mean) |

*GLCM-related features were computed on the three orthogonal planes (XY, XZ, and YZ) passing through the nucleus centroid and then summarized across planes (mean, standard deviation, minimum, maximum, and median).

**Extended Data Table 2.** A list of 3D supercell graphs features.

| Global graph topology (4) | Supercell-subtypes interaction frequencies (\|S\| × S\|) | Neighborhood enrichment (z-scores) (\|S\| × \|S\|) | Cell-type centrality scores (\|S\|× 3) |
|---|---|---|---|
| node_count, graph_density, graph_avg_degree, graph_clustering_coeff | inter_freq_{$S_i$}_{$S_j$} | enrichZ_{$S_i$}_{$S_j$} | Degree_centrality (\|S\|), Average_clustering (\|S\|), Closeness_centrality (\|S\|), |

*S: number of supercell subtypes; $S_i$: subtype i; $i, j \in \{1, \ldots, S\}$

**Extended Data Table 3.** Statistical comparison of top 10 highly stable features between non-BCR and BCR groups.

| Feature | U_statistic | p_value | p_adj | Direction |
|---|---|---|---|---|
| cancer_lumen_mean_dist2mass | 1017 | 0.001 | 0.0034 | Higher in non-BCR |
| std_enrichZ_E2_E2 | 329 | 8.148e-05 | 4.074e-04 | Higher in BCR |
| cancer_stroma_std_dist2mass | 545 | 0.0954 | 0.0954 | Higher in BCR |
| stroma_std_cyto_max_intensity | 888 | 0.0535 | 0.0669 | Higher in non-BCR |
| std_enrichZ_E1_E2 | 254 | 2.256e-06 | 2.256e-05 | Higher in BCR |
| mean_enrichZ_E2_S3 | 991 | 0.0026 | 0.0064 | Higher in non-BCR |
| median_E3_closeness_centrality | 948 | 0.0104 | 0.0174 | Higher in non-BCR |
| std_enrichZ_E2_S3 | 426 | 0.0035 | 0.007 | Higher in BCR |
| max_inter_freq_E3_S1 | 870.5 | 0.0807 | 0.0897 | Higher in non-BCR |
| stroma_mean_cyto_std_intensity | 940 | 0.0132 | 0.0189 | Higher in non-BCR |

Each feature distributions were compared between patients without BCR within 5 years after surgery (non-BCR, n = 44) and patients with BCR within 5 years after surgery (BCR, n = 32). Group differences were assessed using two-sided Mann–Whitney U tests, with the U statistic and exact P value reported for each feature. P values were adjusted across the tested features using the Benjamini–Hochberg false discovery rate procedure.

**Extended Data Table 4.** NMI across a range of supercell formation parameters (r=10-30 μm).

| Supercell | Mean NMI |
|---|---|
| Supercells_r10 | 0.68 |
| Supercells_r15 | 0.68 |
| Supercells_r20 | 0.70 |
| Supercells_r25 | 0.66 |
| Supercells_r30 | 0.65 |

**Extended Data Table 5.** Statistical comparison of supercell- and cell-level NMI stability .Statistical significance was assessed using two-sided paired t-tests comparing supercell-level and cell-level NMI values across 10 replicates.

| Feature noise $\alpha$ | Supercell configuration | t_statistic | df | p_value | p_adj | shapiro_p |
|---|---|---|---|---|---|---|
| 0.05 | Supercells_r10 | 11.96 | 9 | 7.92e-07 | 2.97e-06 | 0.34 |
| 0.05 | Supercells_r15 | 9.11 | 9 | 7.73e-06 | 1.05e-05 | 0.86 |
| 0.05 | Supercells_r20 | 12.747 | 9 | 4.60e-07 | 2.39e-06 | 0.16 |
| 0.05 | Supercells_r25 | 9.65 | 9 | 4.81e-06 | 8.02e-06 | 0.19 |
| 0.05 | Supercells_r30 | 9.708 | 9 | 4.58e-06 | 8.02e-06 | 0.14 |
| 0.1 | Supercells_r10 | 10.494 | 9 | 2.39e-06 | 5.98e-06 | 0.87 |
| 0.1 | Supercells_r15 | 9.467 | 9 | 5.64e-06 | 8.45e-06 | 0.6 |
| 0.1 | Supercells_r20 | 10.693 | 9 | 2.04e-06 | 5.98e-06 | 0.21 |
| 0.1 | Supercells_r25 | 7.007 | 9 | 6.28e-05 | 6.73e-05 | 0.7 |
| 0.1 | Supercells_r30 | 7.835 | 9 | 2.61e-05 | 3.02e-05 | 0.77 |
| 0.2 | Supercells_r10 | 16.489 | 9 | 4.95e-08 | 7.42e-07 | 0.59 |
| 0.2 | Supercells_r15 | 12.687 | 9 | 4.78e-07 | 2.39e-06 | 0.5 |
| 0.2 | Supercells_r20 | 9.929 | 9 | 3.80e-06 | 8.02e-06 | 0.77 |
| 0.2 | Supercells_r25 | 7.866 | 9 | 2.53e-05 | 3.02e-05 | 0.92 |
| 0.2 | Supercells_r30 | 6.072 | 9 | 1.85e-04 | 1.85e-04 | 0.52 |

Statistical significance was assessed using two-sided paired t-tests. P values were adjusted across all comparisons using the Benjamini–Hochberg false discovery rate procedure. Shapiro–Wilk tests were used to assess the normality of paired differences before applying paired t-tests.

**Supplementary Video 1 .** Video abstract summarizing the SCALE3D study workflow and major findings.

**Supplementary Video 2.** Three-dimensional visualization of the SCALE3D workflow.

Videos are provided as separate files
